# Attractive and repulsive couplings between circadian pacemaker neurons promote entrainment to light-dark cycles with after-effect

**DOI:** 10.1101/2025.11.20.689452

**Authors:** Yuta Kitaguchi, Satoru Komori, Hajime Tei, Koichiro Uriu

## Abstract

The mammalian circadian pacemaker, suprachiasmatic nucleus (SCN), comprises a core region that receives light signals from the retina and a shell region that outputs pacemaking rhythms to peripheral tissues. The free-running period (FRP) of animals in constant darkness (DD) correlates with the period of the preceding LD cycle, a phenomenon known as after-effect. To elucidate the mechanisms underlying robust entrainment and after-effect, we analyzed phase oscillator models incorporating attractive and repulsive couplings that respectively decrease or increase phase differences between oscillators. Attractive coupling from the core to shell regions accounts for experimentally observed phase relationships between these regions under LD cycles. Remarkably, repulsive coupling from the shell to core regions promotes SCN entrainability to LD cycles. Furthermore, after transfer to DD, the FRP slowly returns to the intrinsic SCN period, reflecting the after-effect. Our analysis identifies an SCN coupling architecture that underlies stable daily activity rhythms.

## Introduction

Organisms experience seasonal changes in day–night length throughout the year. Such variations must be internalized by plants [1] and animals [2] through synchronization of their internal circadian rhythms with external light cycles, a process termed entrainment. A central question is how the circadian clock achieves stable entrainment to a light–dark (LD) cycle. A classical paradigm for studying this mechanism is to entrain the circadian clock to non-24-hour LD cycles referred to as T-cycles [3].

In mammals, the suprachiasmatic nucleus (SCN) of the brain acts as the central pacemaker of the circadian clock [4, 5]. Oscillations in the electrophysiological activity of SCN neurons are required to maintain circadian rhythmicity under constant conditions [6], and arise from the circadian expression of clock genes [7]. Among these, delayed negative feedback regulation of *Period1/2* (*Per1/2*) genes is responsible for the generation of autonomous transcriptional rhythms [7]. The SCN forms a complex network of approximately 10^4^ neurons, divided into ventrolateral core and dorsomedial shell regions [5, 8–10]. The core region contains neurons secreting vasoactive intestinal peptide (VIP), whereas the shell region contains neurons secreting arginine-vasopressin (AVP) [8, 11]. VIP release and AVP production are regulated by neuronal firing activity and circadian clock proteins, respectively [12–14]. These neurons also express both VIP and AVP receptors [15–17], receiving phase information from neighboring SCN neurons. Intercellular coupling between VIP and AVP neurons synchronizes circadian gene expression by reducing phase differences between them (attractive coupling) [14, 18–23]. At the same time, most SCN neurons are GABAergic [8], and GABA signaling tends to increase phase differences between oscillators (repulsive coupling) [24, 25]. Indeed, GABA-mediated coupling can decrease synchronization among SCN neurons [26]. Consequently, the collective circadian rhythm of the SCN emerges from the combined effects of attractive and repulsive couplings.

In addition to generating autonomous rhythms, the SCN also responds to light. Core neurons receive photic input from the retina via the retinohypothalamic tract [5]. In these neurons, light-induced transcription of *Per1/2* advances or delays circadian gene expression rhythms depending on stimulus timing [27–31]. Then, the information of the phase shift in the core neurons is transmitted to the shell region primarily through VIP [8,18,23,32], causing the phase shift of the shell neurons. Finally, the shell neurons output circadian signals that entrain peripheral clocks, completing the phase shift at an individual level. Thus, the mammalian behavioral rhythms can entrain to LD cycles with near 24-hour periodicity.

When rodents are entrained to a non-24-hour LD cycle (T-cycle) and subsequently transferred to constant darkness (DD), their free-running period (FRP) in DD reflects that of the previous cycle [3, 33– 35]. Mice entrained to an 11-hour light/11-hour dark (LD 11:11, T22) showed a shorter FRP than those entrained to a T24 (LD 12:12) [33]. In contrast, the FRP after T26 (LD 13:13) was longer than that after T24. The FRP after non-24 T-cycles slowly returned to a species-specific period with a characteristic time of 20 ∼ 47 days (Fig. 1) [3, 35]. This after-effect, observed across mammals, birds and insects [3], strongly suggests that the circadian clock retains information about the period of previously experienced LD cycles.

**Figure 1.**
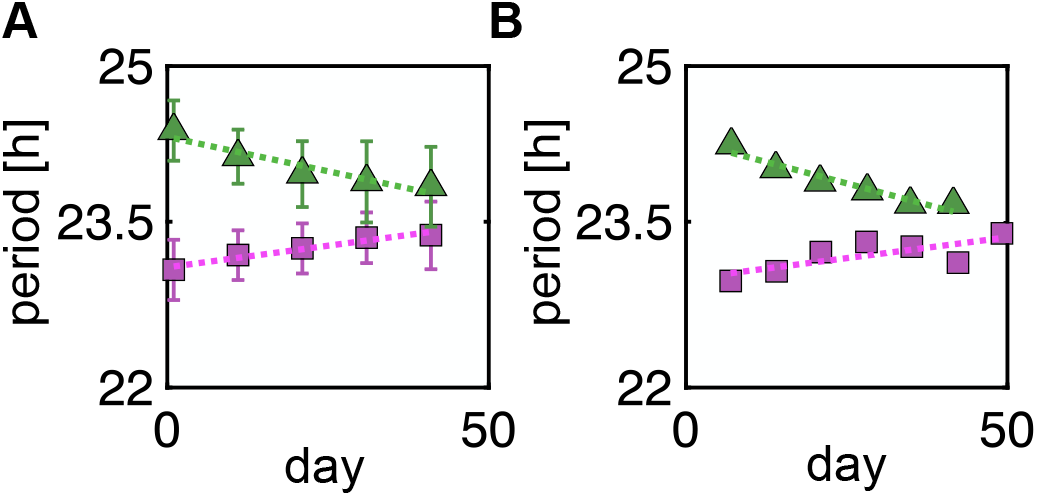
Changes in mouse FRPs in DD after non-24 T-cycles. (A), (B) Timeseries of average FRP from (A) Pittendrigh et al. [3] (*n* = 14 for T20; *n* = 12 for T28) and (B) Aton et al. [35] (*n* = 9 for T20; *n* = 15 for T28). Magenta squares represent average FRPs after T20 (LD 10:10) and green triangles represent those after T28 (LD 14:14). Dotted lines indicate exponential fit *e*^*t/ξ*^ to the data with a characteristic time *ξ*. We obtained *ξ* = 47 days for T20 in (A), *ξ* = 37 days for T20 in (B), *ξ* = 27 days for T28 in (A) and *ξ* = 21 days for T28 in (B). In (A), we used all the data points from the mouse population shown in Fig. 8 of Pittendrigh and Daan [3]. In (B), we used the data from mice younger than 3 months of age presented in Fig. 5 of Aton et al. [35]. The horizontal axis (day) is shown on a linear scale, and the vertical axis (period) on a logarithmic scale. Error bars represent standard deviations. Data extraction and model fitting are described in Methods.

Given the SCN’s role as the central pacemaker, the after-effect in behavioral rhythms likely reflects changes in SCN state induced by non-24 T-cycles [34–36]. Monitoring *Per1/2* transcriptional cycling in SCN slices from mice entrained to non-24 T-cycles revealed phase differences between the core and shell regions [34]. In mice entrained to T24, the phase of oscillation in the SCN is spatially uniform, showing no phase difference between the two regions. In contrast, in mice entrained to T22, the core phase advanced by 2 hours relative to shell. In mice entrained to T26, it was delayed by 2 hours. Importantly, whether these phase differences contribute to the entrainment and the after-effect remains unclear.

Previous theoretical studies have described the after-effect using coupled oscillator models [34, 37, 38]. Daan and Berde (1978) showed that it can arise in a two-oscillator system describing evening (E) and morning (M) oscillators that govern activity onset and offset in rodents [38]. They showed that non-24 T-cycles modulate the phase difference between the E and M oscillators, and that the after-effect appears as a transient dynamics as this difference gradually returns to its steady state value in DD [38]. Azzi et al. (2017) experimentally observed changes in DNA methylation of genes related to neurotransmitter receptors and ion channels –including potassium, calcium and GABA channels –in the SCN, depending on the period of the experienced non-24 T-cycle [34]. Based on these observations, they proposed that non-24 T-cycles alter the balance between attractive and repulsive couplings from the shell to core regions, resulting in a positive correlation between FRP and the experienced T-cycle period [34]. Recently, a mathematical model incorporating neuronal synaptic plasticity was examined [37]. This model assumed that SCN neurons with intrinsic periods close to a non-24 T-cycle exert stronger influence over other oscillators, enabling entrainment of the SCN to the LD cycle. The observed DNA methylation and synaptic plasticity likely contribute to the after-effect by forming cellular-level memory of prior LD cycles. However, previous mathematical models did not examine how phase differences between the SCN core and shell regions induced by non-24 T-cycles [34] contribute to the after-effect. Moreover, despite extensive studies, the biological significance of the after-effect has long been unknown.

By extending previous modeling work, this study connects (1) the mechanism of robust SCN entrainment to non-24 T-cycles, and (2) the mechanism by which the phase difference between the core and shell regions alter the SCN period in DD and prolong the after-effect. We first analyze a two-oscillator system that describes the SCN core and shell regions coupled to an LD cycle. Then, we examine the after-effect in a model that describes the circadian rhythms of multiple SCN neurons as a coupled oscillator network.

We show that the enhanced SCN entrainability through the combined action of attractive and repulsive couplings naturally gives rise to a prolonged after-effect.

## Results

### Phase difference formation by T-cycles

Because the behavioral period of mammals is primarily regulated by the SCN [4], the after-effect can be attributed to the slow recovery of the integrated circadian period in the SCN. To describe the changes in the SCN during the after-effect, we model interactions between the circadian clocks in the core and shell regions. The core region receives photic input from the retina via the retinohypothalamic tract and transmits this information to the shell region [5, 9]. This SCN architecture has been modeled in previous work using coupled phase oscillators [25,34,39,40]. Based on these works, we first consider a two-oscillator model of the SCN where *ϕ*_*C*_ represents the phase of the core region and *ϕ*_*S*_ that of the shell region, coupled to the phase of a LD cycle *ϕ*_*L*_ (Fig. 2A) :

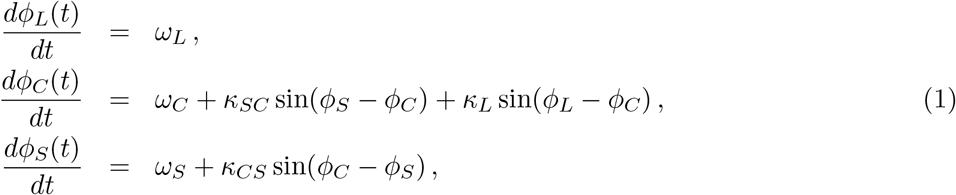

where *ω*_*L*_ is the frequency of the LD cycle specifying T-cycle period *T* = 2*π/ω*_*L*_, and *ω*_*C*_ and *ω*_*S*_ are the intrinsic frequencies of the core and shell regions, respectively. For the illustration of the mechanism of after-effect, we first consider identical intrinsic frequencies *ω*_*C*_ = *ω*_*S*_ = *ω*. In a later section, we analyze the effect of frequency difference *ω*_*C*_ *≠ ω*_*S*_. For simplicity, we assume a sinusoidal coupling function between the core, shell, and light phases in Eq. (1). The coupling between these oscillators depends on their phase differences. *κ*_*SC*_ and *κ*_*CS*_ are the coupling strengths from shell to core and from core to shell, respectively. These coupling strengths can be either positive or negative. A positive value represents attractive coupling that reduces phase difference between oscillators, whereas a negative value represents repulsive coupling that increases it. Unlike a previous mathematical model [34] where coupling strength varied depending on the T-cycle period, here we fix the values of *κ*_*SC*_ and *κ*_*CS*_ for different T-cycle periods. *κ*_*L*_ is the coupling strength of the core phase to the LD phase. As we will show below, the value of *κ*_*L*_ does not affect phase difference formation between the core and shell regions under a T-cycle. We therefore assume a positive coupling strength *κ*_*L*_ = 1 throughout this paper unless mentioned otherwise.

**Figure 2.**
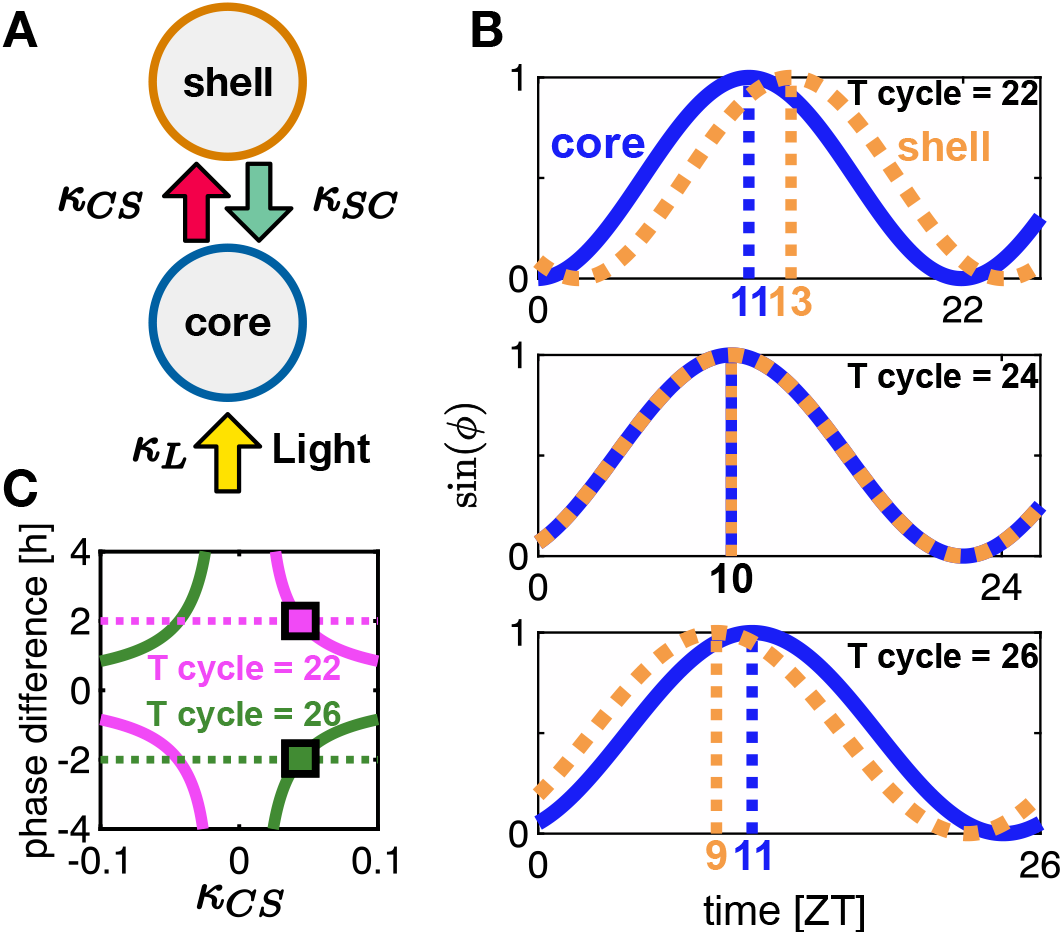
Phase difference formation between the core and shelll regions of the SCN by non-24 T-cycles. (A) Two-oscillator model. The SCN is divided into the core and shell regions, with the core region receiving light input. *κ*_*L*_ is the coupling strength from the light phase to the core phase. Received light information is transferred to the shell region with the coupling strength *κ*_*CS*_ . The shell region feeds back the circadian signal to the core region with the coupling strength *κ*_*SC*_ . (B) The phase difference between the core and shell regions entrained to T22, T24 and T26 is illustrated using sin(*ϕ*) as a function of zeitgeber time (ZT). The zeitgeber times *t*_*Z*_ satisfying *ϕ*(*t*_*Z*_ ) = *π/*2 (sin *ϕ* = 1) for the core and shell regions are highlighted with dotted vertical lines. These were determined based on the acrophases of experimental PER2 oscillation data [34]. (C) Estimation of the coupling strength *κ*_*CS*_ by fitting the phase difference *ψ*_*CS*_ to previous experimental data [34]. Dotted horizontal lines indicate experimentally obtained phase differences *ψ*_*CS*_ between the core and shell regions. Solid lines are relationships between *ψ*_*CS*_ and *κ*_*CS*_ obtained from Eq. (2). Magenta and green lines indicate results for T22 and T26, respectively. Squares at *κ*_*CS*_ = 0.045 indicate matches between model estimation and experimental data.

The core and shell phases can be entrained to a LD cycle with non-24-hour period (Fig. 2B). By analytical calculation described in Supplementary Information, Text S1, we obtain the phase difference between the core and shell regions *ψ*_*CS*_ = *ϕ*_*C*_ − *ϕ*_*S*_ in T-cycle as:

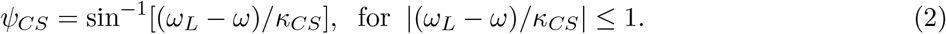

*ψ*_*CS*_ in Eq. (2) depends on the frequency mismatch between T-cycle and SCN, and coupling strength from core to shell *κ*_*CS*_ (Fig. 2B, C). However, it is independent of coupling from shell to core *κ*_*SC*_. The FRP of behavioral rhythms of wildtype mice is usually longer than 22 hours (*ω*_*T* 22_ − *ω >* 0) and shorter than 26 hours (*ω*_*T* 26_ − *ω <* 0) [33, 34]. Therefore, experimentally observed phase differences between the core and shell regions under these T-cycles [34] can be reproduced if

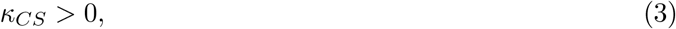

suggesting an attractive coupling from the core to shell regions (Fig. 2B, C). Figure 2C also indicates that *ψ*_*CS*_ becomes smaller as the magnitude of *κ*_*CS*_ increases. We obtain a quantitative agreement between experimental and simulation data if *κ*_*CS*_ ≈ 0.05 h^−1^ (Fig. 2C; Methods). Both experimental and simulation data provide 2 and −2 hours phase difference for T22 and T26, respectively [34]. Although Fig. 2C shows other intersection points between the dotted and solid lines in the region *κ*_*CS*_ *<* 0, the phase relationship between the core and shell regions at these points contradicts the experimental observations, since the core phase is behind (ahead of) the shell phase in T22 (T26).

Mice were also successfully entrained to T20 and T28 in other previous studies [3, 35]. We infer the phase differences between the core and shell regions of these mice under T20 and T28 [3, 35] using this estimated value of *κ*_*CS*_. The SCN with the intrinsic period of 24 h can entrain to T28 with this value of *κ*_*CS*_, and the phase of shell region advances by 3.7 hours from that of the core region. In contrast, T20 is out of the entrainment range for the shell region Eq. (2) with this value of *κ*_*CS*_. However, a slight increase in the value of *κ*_*CS*_ (*κ*_*CS*_ ≥ 0.0524) is required for the entrainment, and shell phase is delayed 5.9 h relative to the core phase.

### Entrainment range for T-cycles

The entrainment condition of the shell region to a non-24 T-cycle is given by the inequality in Eq. (2). After similar calculations, we obtain the entrainment condition of the core region to a non-24 T-cycle (Supplementary Information, Text S2):

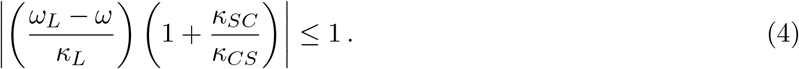

Inequality (4) indicates that the entrainment range can be extended by the coupling strength *κ*_*L*_. To visualize this relationship, we plot the entrainment range as a purple shade in a *ω*_*L*_ − *ω* (frequency mismatch) and *κ*_*L*_ parameter space in Fig. 3A, B. The shape of the entrainment range is referred to as an Arnold tongue [41]. The condition (4) also suggests that the width of the Arnold tongue extends if −1 *< κ*_*SC*_*/κ*_*CS*_ *<* 0 as (1 + *κ*_*SC*_*/κ*_*CS*_) becomes smaller (Fig. 3B). Because *κ*_*CS*_ *>* 0 in Eq. (2), this leads to:

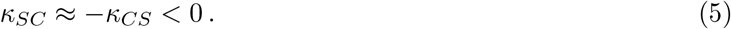

**Figure 3.**
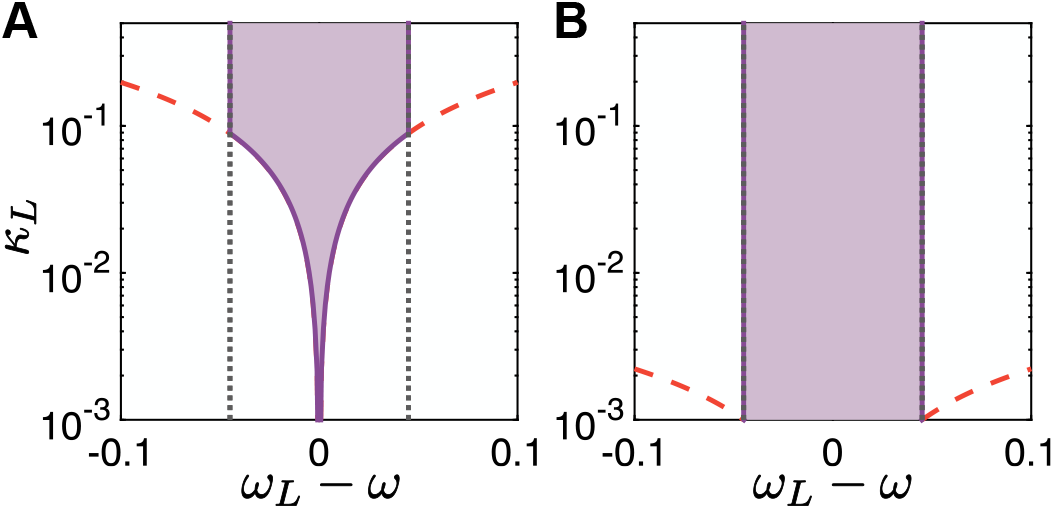
Enlargement of the entrainment range by repulsive coupling from the shell to core regions. (A), (B) Arnold tongue showing the dependence of entrainment on frequency mismatch *ω*_*L*_ − *ω* and coupling strength from the core region to LD cycles *κ*_*L*_. (A) *κ*_*SC*_ = 0.045. (B) *κ*_*SC*_ = −0.045. The horizontal axis is shown on a linear scale and the vertical axis on a logarithmic scale. Black dotted lines indicate the condition Eq. (2), and red dashed lines indicate the condition Eq. (4). The purple shaded area indicates entrainment of both the core and shell regions to LD cycles.

Thus, a combination of a repulsive coupling from the shell to core regions and an attractive coupling from the core to shell regions promotes the entrainment of the SCN to LD cycles. An intuitive explanation of how repulsive coupling enhances light entrainment is as follows. A non-24 T-cycle induces a phase difference between the core and shell regions (Fig. 2). Because the core region directly receives light input while the shell region does not, the phase difference between the core and LD phases is smaller than that between the shell and LD phases. In other words, the core phase lies between the LD phase and the shell phase. With an attractive coupling from the shell to the core regions, the shell phase pulls the core phase away from the LD phase. Therefore, a large *κ*_*L*_ is required for the core phase to entrain to the LD phase (Fig. 3A). In contrast, a repulsive coupling causes the shell phase to push the core phase toward the LD phase, thereby reducing the *κ*_*L*_ value required for entrainment (Fig. 3B).

### After-effect

Next, we study the relationship between the phase difference *ψ*_*CS*_ and the after-effect in DD. To model constant darkness after a non-24 T-cycle, we set *κ*_*L*_ = 0 in Eq. (1) in the following analysis. Since we determined the value of *κ*_*CS*_ from experimental data [34] as described above, we analyze the period change in DD by varying the coupling strength from the shell to core regions *κ*_*SC*_ (Methods).

If *κ*_*SC*_ is positive, the periods of the core and shell regions quickly return to their intrinsic values 2*π/ω* in DD (Fig. S1). However, if *κ*_*SC*_ is negative and its magnitude is close to *κ*_*CS*_, we observe a slow period change in DD after a non-24 T-cycle (Fig. 4A, B). After T22, the period in DD remains shorter than the intrinsic period 2*π/ω* = 24 for more than two months. In contrast, the period in DD remains longer after T26. The phase difference between core and shell regions *ψ*_*CS*_ also gradually decreases at the same rate as the period changes (Fig. 4B). The half-life of the period increases as *κ*_*SC*_*/κ*_*CS*_ approaches −1 (Fig. 4C). This half-life is independent of the initial phase difference in DD (Fig. S2).

**Figure 4.**
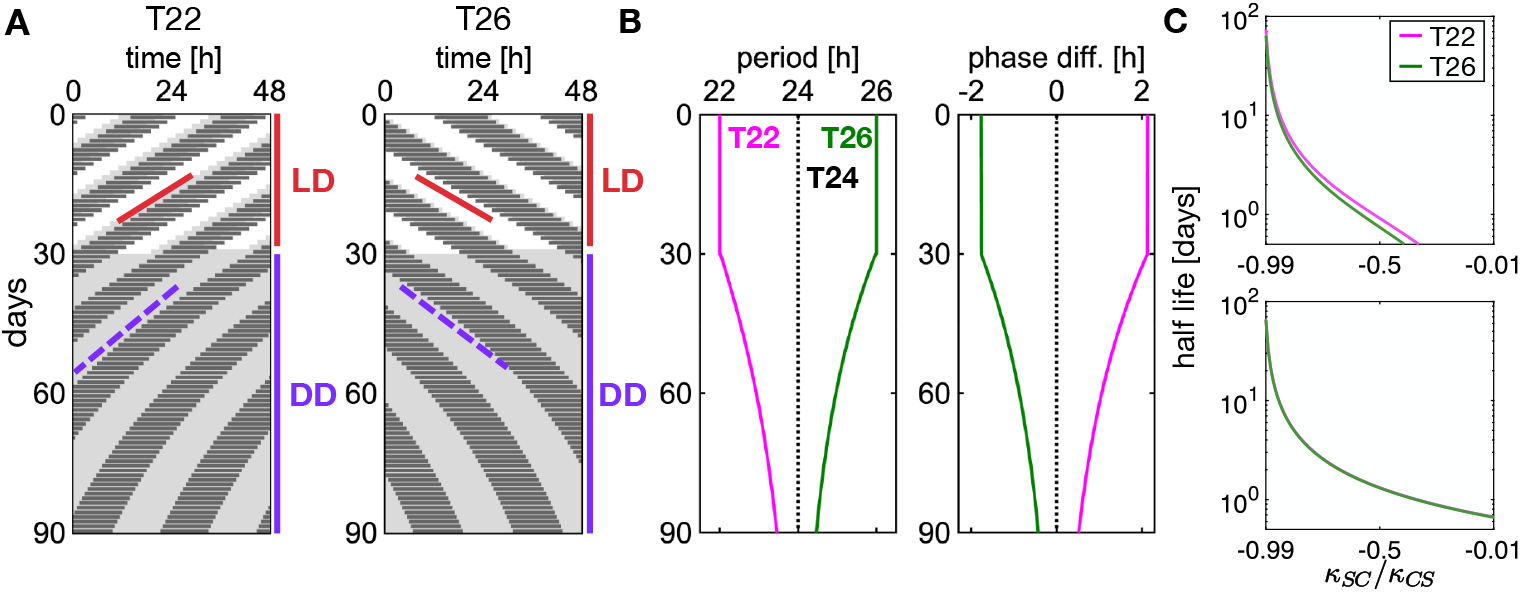
After-effect by the combination of attractive and repulsive couplings between the core and shell regions. (A) Simulated double-plot actograms under non-24 T-cycles (T22: left, T26: right) and subsequent DD. Black horizontal bars represent the active phases of animals defined by the shell phase *ϕ*_*S*_ in the interval between *π* and 2*π*. Solid red and broken purple slopes indicate the activity onsets defined as *ϕ*_*S*_ = *π*. (B) Timeseries of (left) collective period of core and shell 2*π/*Ω and (right) their phase difference *ψ*_*CS*_ . Ω is the collective frequency 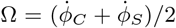. Simulation results under LD cycles from day 0 to 30 (red vertical lines in A) and under DD from day 30 to 90 (purple vertical lines in A). To represent the SCN phase difference *ψ*_*CS*_ by a unit of time (hours), we divide *ψ*_*CS*_ by 2*π/*24 in both a LD cycle and DD for continuity of phase difference at the transition at day 30 (from LD to DD). (C) Dependence of the half-life of FRP (top) and phase difference (bottom) on the ratio *κ*_*SC*_ */κ*_*CS*_ . *κ*_*CS*_ is fixed as shown in Table 1 while *κ*_*SC*_ is varied within the range |*κ*_*SC*_ | ≤ *κ*_*CS*_ . In (B) and (C), magenta and green lines represent results in DD following entrainment to T22 and T26, respectively.

To derive the mathematical relationship between the period and the phase difference, we first define a collective frequency of the core and shell regions 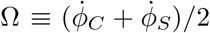 where 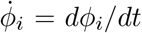. The collective period is denoted as

**Table 1.**
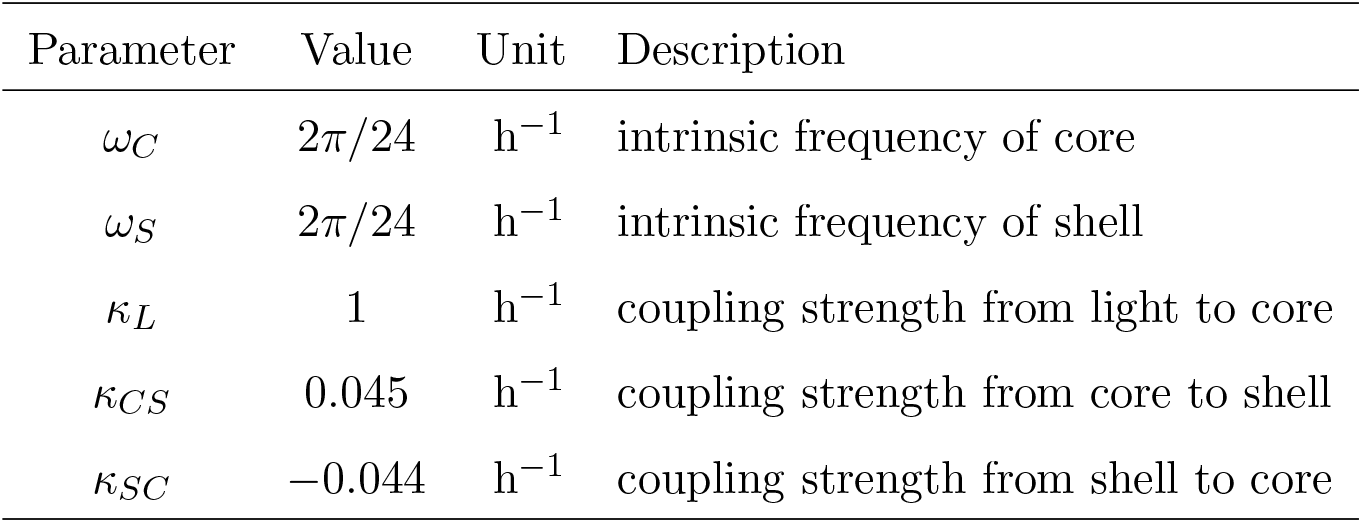
Parameter values for the numerical simulation of the two-oscillator model.

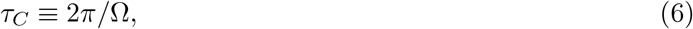

which represents a FRP in DD. We obtain the collective frequency in DD as a function of the phase difference (Supplementary Information, Text S3)

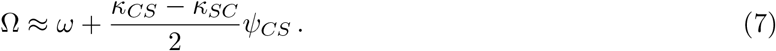

The first term of Eq. (7) is the intrinsic frequency of the two SCN subregions. The second term is the contribution of the phase difference *ψ*_*CS*_ to the collective frequency. Eq. (7) indicates that the collective frequency is proportional to the phase difference if

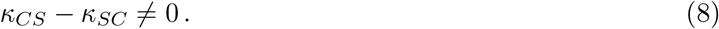

Next, we analyze the dynamics of the phase difference *ψ*_*CS*_ in DD. If |*ψ*_*CS*_| is small, we may approximate its time derivative as:

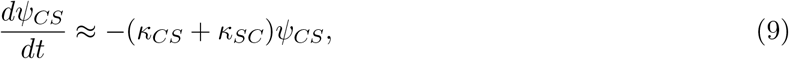

see Supplementary Information, Text S3 for the derivation. The solution of this linear differential equation is:

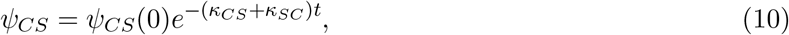

where *ψ*_*CS*_(0) is the phase difference at the onset of DD. This phase difference decays if:

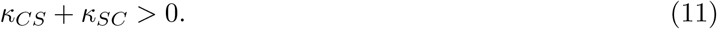

We define the half-life of the phase difference as 1*/τ*_*h*_ ≡ *κ*_*CS*_ + *κ*_*SC*_. Since we determined that *κ*_*CS*_ *>* 0, the half-life becomes long if

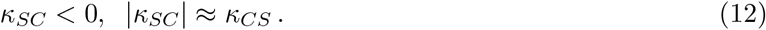

Eq. (8) also suggests that *ψ*_*CS*_ affects Ω with *κ*_*SC*_ *<* 0. We write *κ*_*SC*_ = −|*κ*_*SC*_| in Eq. (7) and obtain

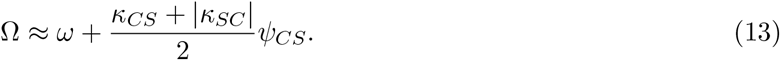

After T22, the phase difference between the core and shell regions *ψ*_*CS*_ is positive (Fig. 2B), indicating Ω *> ω* and 2*π/*Ω *<* 2*π/ω*. After T26, *ψ*_*CS*_ is negative (Fig. 2B), and 2*π/*Ω *>* 2*π/ω*. Importantly, condition (12) corresponds to that for enhanced SCN entrainability to LD cycles Eq. (5). Thus, the combination of attractive and repulsive couplings enhances the entrainment of the SCN to LD cycles accompanied by a positive correlation between the period of non-24 T-cycles and the subsequent FRP in DD. This combination also produces a gradual reduction in the phase difference between the core and shell regions, resulting in a prolonged after-effect.

### Difference in intrinsic frequency between the core and shell regions

In the previous section, we studied the after-effect by assuming the same intrinsic frequency of the core and shell regions *ω*_*C*_ = *ω*_*S*_ = *ω* for simplicity. However, desynchronization experiments of the rat SCN using chemical and physical perturbations demonstrated that the dorsomedial region oscillates at a higer frequency than the rest of the SCN [42]. Dissection of the dorsal and ventral regions in the mouse SCN revealed that the frequency of the dorsal region is higher than that of the ventral region [43]. Coupled oscillator models incorporating intrinsic frequency differences between SCN subregions reproduced experimentally observed spatial phase waves propagating from the medial to lateral SCN regions [42–44]. Motivated by these experimental and theoretical findings, we next investigate whether the intrinsic frequency difference between the core and shell regions (*ω*_*C*_ ≠*ω*_*S*_) influences the after-effect. If this is the case, we may infer the intrinsic frequency difference in living animals from the observation of the after-effect.

In the presence of frequency difference Δ*ω* = *ω*_*C*_ − *ω*_*S*_, the steady state of phase difference 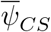 in DD is not necessarily equal to zero (Supplementary Information, Text S4). Such non-zero phase difference makes it difficult to compare results obtained using different values of Δ*ω*. Hence, we introduce a phase shift parameter *α* and tune its value to keep 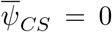 in DD for better comparison (Supplementary Information, Text S4):

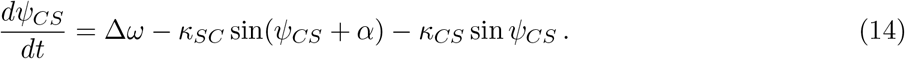

We find that the difference in intrinsic frequency Δ*ω* generates a T-cycle dependent relaxation of FRP (Fig. 5A-C). When Δ*ω <* 0, the changes in FRP and phase difference are slower after T26 than T22 (Fig. 5A, C). Conversely, when Δ*ω >* 0, the changes are slower after T22 than T26 (Fig. 5B, C). The half-life of the collective period in DD increases with Δ*ω* after T22 (magenta line in Fig. 5C) while it decreases with Δ*ω* after T26 (green line).

**Figure 5.**
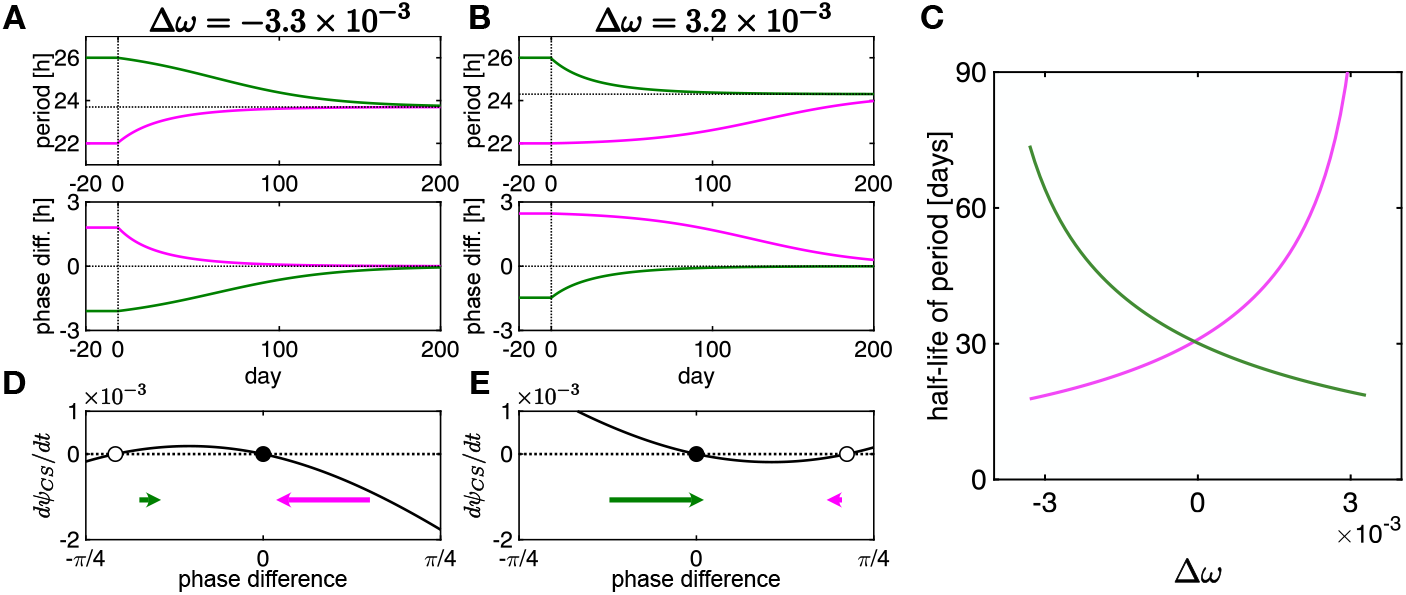
After-effect in the presence of the intrinsic frequency difference Δ*ω* = *ω*_*C*_ − *ω*_*S*_ between the core and shell regions. (A), (B) Timeseries of (top) the collective period and (bottom) the phase difference between the core and shell phases for (A) Δ*ω* = −3.3 *×* 10^−3^ and (B) Δ*ω* = 3.2 *×* 10^−3^. The core and shell regions are entrained to a non-24 T-cycle from day -20 to day 0, and released to DD from day 0 (dotted vertical lines) in the simulations. Magenta and green lines show the timeseries after T22 and T26, respectively. (C) Dependence of the half-life of the collective period 2*π/*Ω in DD on Δ*ω* after T22 (magenta) and T26 (green). (D), (E) *dψ*_*CS*_ */dt* as a function of phase difference *ψ*_*CS*_ . The green and magenta arrows indicate the flow of *ψ*_*CS*_ . The length of the arrows represents the magnitude |*dψ*_*CS*_ */dt*| in each region. Filled and open circles represent stable and unstable steady state phase differences, respectively.

We linearize Eq. (14) to seek a simpler description of phase difference decay, but such linear analysis does not reveal the mechanism for asymmetrical after-effect. To show the mechanism in the non-linear regime, we plot *dψ*_*CS*_*/dt* in Eq. (14) as a function of *ψ*_*CS*_ in Fig. 5D, E. In Fig. 5D, E, intersections at *dψ*_*CS*_*/dt* = 0 represent steady state phase differences between the core and shell regions. The steady state 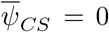 is stable (filled circles). In addition, there is an unstable steady state phase difference 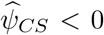 (open circles). When the intrinsic frequency of the core region is lower than that of the shell region Δ*ω <* 0 as in Fig. 5D, *dψ*_*CS*_*/dt >* 0 in 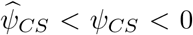 and the speed |*dψ*_*CS*_*/dt*| is smaller than in *ψ*_*CS*_ *>* 0. Therefore, the positive *ψ*_*CS*_ caused by T22 quickly returns to the stable steady state, while the negative *ψ*_*CS*_ caused by T26 returns much more slowly (Fig. 5A, D). In contrast, when Δ*ω >* 0, an unstable steady state is 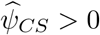 (Fig. 5E). In this case, the speed |*dψ*_*CS*_*/dt*| is smaller in 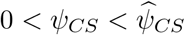 than in *ψ*_*CS*_ *<* 0, causing slower decrease of phase difference after T22 than T26 (Fig. 5B, E).

We also solve the nonlinear differential equation (14) analytically (Supplementary Information, Text S5) [45, 46]. The exact solution indicates that the prolonged after-effect occurs if

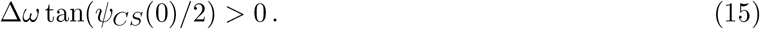

Condition Eq. (15) holds after T26 with Δ*ω <* 0 (Fig. 5A) and after T22 with Δ*ω >* 0 (Fig. 5B). Intuitively, when the phase of the faster oscillator leads that of the slower one, it requires more time for the phase difference to decrease. For instance, after T22 with Δ*ω* = *ω*_*C*_ − *ω*_*S*_ *>* 0, the phase of the faster core regions is ahead of that of the slower shell region *ψ*_*CS*_(0) *>* 0, so the shell region requires a longer time to catch up with the core region.

The asymmetric after-effect shown in Fig. 5 has not yet been reported in mammals. Our theoretical result predicts that if animals released from T22 and T26 show different after-effect durations, the intrinsic frequencies between the core and shell regions may also differ.

### Entrainment and after-effect in the complex neuronal network

So far, we have examined entrainment and the after-effect using the two-oscillator system Eq. (1), which represents the phases of the core and shell regions of the SCN coupled to a non-24 T-cycle. Next, we construct a mathematical model describing the phase dynamics of individual SCN neurons to validate the results obtained from the simplified two-oscillator system within a more complex and detailed neural network.

We develop an SCN network model comprising *N*_*C*_ core neurons and *N*_*S*_ shell neurons (Fig. 6A and Supplementary Information, Text S6) [47]. The total number of neurons is *N* = *N*_*C*_ + *N*_*S*_ and we assume *N*_*C*_ ≤ *N*_*S*_ based on experimental observations [14]. We use both directed and undirected graphs to describe the interaction network between neurons. Each of the core and shell regions is described by a two-dimensional lattice (Fig. 6A). Neuron *i* (1 ≤ *i* ≤ *N*_*C*_ for a core neuron and *N*_*C*_ + 1 ≤ *i* ≤ *N*_*C*_ + *N*_*S*_ for a shell neuron) mutually interacts (i.e., undirected) with its four nearest neighbors and with 12 additional distant neurons belonging to the same SCN subregion (core or shell) as neuron *i*. For these 12 distant couplings within each SCN subregion, the nearest neighbors are excluded, and 12 target cells are randomly selected regardless of distance. We assume an attractive coupling between neurons of the same subregion, *κ*_*CC*_ *>* 0 and *κ*_*SS*_ *>* 0 for the core and shell regions, respectively. For coupling between core and shell neurons, we assume that *N*_*CS*_ core neurons send directed coupling to shell neurons (Fig. 6A, B). Each shell neuron receives a single connection from only one core neuron. *N*_*SC*_ shell neurons send directed connections to core neurons. We assume *N*_*SC*_ ≤ *N*_*C*_ so that a single core neuron receives either zero connection or exactly one connection from a single shell neuron (Fig. 6B). As in the two-oscillator model, the coupling from core to shell neurons *κ*_*CS*_ is assumed to be attractive (*κ*_*CS*_ *>* 0), while the coupling from shell to core neurons *κ*_*SC*_ is repulsive (*κ*_*SC*_ *<* 0). Each core neuron is coupled to the T-cycle with the coupling strength *κ*_*L*_. See Supplementary Information, Text S6 for further details regarding the SCN network model.

**Figure 6.**
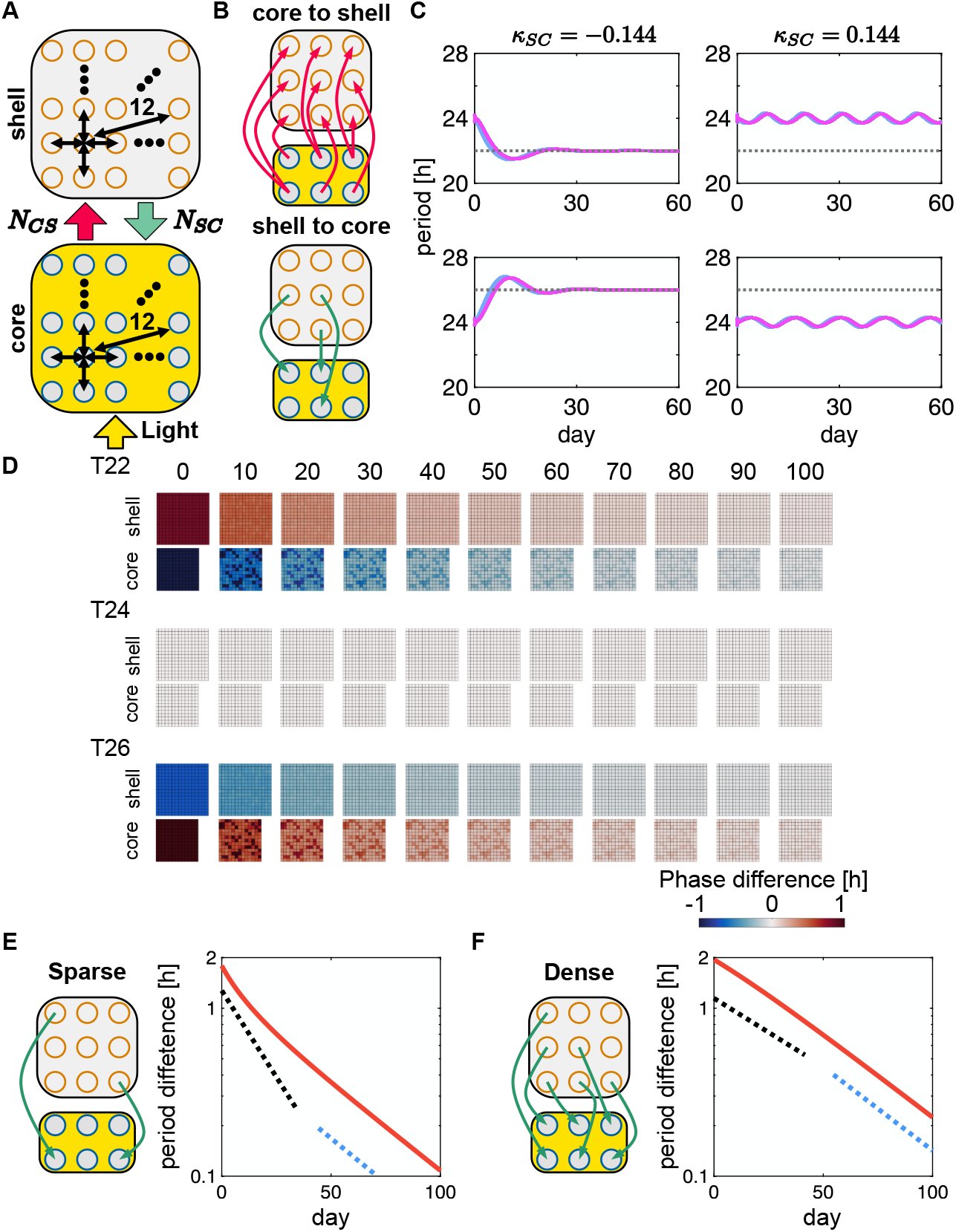
Enhanced entrainability and prolonged after-effect in the SCN network model. (A) Network topology within the core and shell regions. Each neuron is shown as a circle. Based on the actual SCN network, which incorporates diffusive neurotransmitter coupling (local coupling) and synaptic coupling (long-distance coupling), the model includes nearest-neighbor connections and 12 random connections, respectively (black arrows). Core neurons receive directed connections from shell neurons and vice versa, as illustrated in (B). *N*_*CS*_ represents the number of directed connections from core to shell neurons. *N*_*SC*_ represents the number of directed connections from shell to core neurons. All core neurons are coupled to a LD cycle (yellow arrow and background). (B) Directed connections (top) from core to shell neurons and (bottom) from shell to core neurons. (C) Timeseries of the period of SCN neurons under non-24 T-cycles (*χ*=0.23). If the coupling strength from light to core is weak (*κ*_*L*_ = 0.005), SCN neurons with a repulsive coupling from shell to core (*κ*_*SC*_ = −0.144) can entrain to T22 (top left) and T26 (bottom left). In contrast, SCN neurons with an attractive coupling (*κ*_*SC*_ = 0.144) cannot entrain to T22 (top right) and T26 (bottom right) when *κ*_*L*_ = 0.005. Black dotted lines indicate 22 h (top) and 26 h (bottom). The colored lines indicate the periods of individual SCN neurons. (D) Snapshots of phase difference between individual neurons and mean phase from day 0 to 100 (*χ*=0.23). The top, middle, and bottom rows show results in DD after T22, T24, and T26, respectively. (E), (F) Timeseries of period difference in DD for (E) sparse (*χ* = 0.23) and (F) dense (*χ* = 1) directed connections from shell to core neurons. Period difference (vertical axis) is defined as *τ*_*C*_ − 24 where 24 = 2*π/ω* is the intrinsic SCN period. Red lines indicate period differences obtained from numerical simulations. Dotted lines mark the slopes for intervals from day 0 to 10 (black dotted) and from day 95 to 105 (blue dotted). Log-linear plots are used to display exponential changes in period differences.

We examine the effects of repulsive coupling and connectivity from shell to core neurons on entrainability and the after-effect. For this purpose, we define the connectivity:

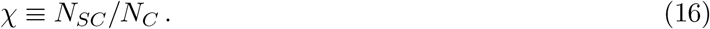

If only a few core neurons receive input from shell neurons, *χ* ≈ 0. A previous study estimated that functional couplings from shell to core neurons are sparse in SCN coronal slices [48]. Conversely, in the case of a dense shell to core connections, *χ* ≈ 1.

We seek to identify the parameter condition for a prolonged after-effect in the neural network model. Analysis of the two-oscillator model showed that the after-effect becomes longer as the coupling ratio *κ*_*SC*_*/κ*_*CS*_ approaches −1 (Fig. 4C). *κ*_*SC*_*/κ*_*CS*_ = −1 corresponds to a bifurcation point in which a steady state phase difference *ψ* = 0 changes its stability as in Eq. (11). In the SCN network model, connectivity *χ* might also affect the stability of *ψ* = 0 as well as *κ*_*SC*_*/κ*_*CS*_. We use a mean-field approximation (Supplementary Information, Text S7) to find the bifurcation point of *ψ* = 0 as an approximation of *κ*_*SC*_ values for long after-effect. We find the bifurcation point as

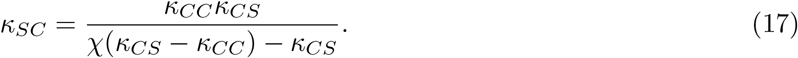

*κ*_*SC*_ values obtained by Eq. (17) agree well with those obtained by numerical linear stability analysis of the original SCN network model (Fig. S3 and Supplementary Information, Text S8). If *κ*_*SC*_ is larger than Eq. (17), the steady state *ψ* = 0 is stable, and phase difference caused by a T-cycle may slowly return to *ψ* = 0. Eq. (17) reveals how *κ*_*SC*_ for a long after-effect depends on coupling from core to shell *κ*_*CS*_ and connectivity *χ*, as well as on coupling within the core region *κ*_*CC*_.

We first simulate the SCN network with a sparse coupling from shell to core, *χ* = 39*/*169 = 0.23 and coupling parameters Eq. (17) (Table 2). We find that under a non-24 T-cycle, the periods of SCN neurons converge to the period of the T-cycle with a repulsive coupling from shell to core neurons *κ*_*SC*_ *<* 0 (Fig. 6C). In contrast, the periods of SCN neurons with an attractive coupling oscillate around 24 h, indicating entrainment failure for the non-24 T-cycles (Fig. 6C). Thus, these results confirm that combining attractive and repulsive coupling promotes entrainment of individual neurons to LD cycles.

**Table 2.** Parameter values for the numerical simulations of the SCN network model.

| Parameter | Value | Unit | Description |
| --- | --- | --- | --- |
| $\omega_C$ | $2\pi/24$ | $\text{h}^{-1}$ | frequency of core |
| $\omega_S$ | $2\pi/24$ | $\text{h}^{-1}$ | frequency of shell |
| $\kappa_L$ | 1 | $\text{h}^{-1}$ | coupling strength from light to core |
| $\kappa_{CS}$ | 0.045 | $\text{h}^{-1}$ | coupling strength from core to shell |
| $\kappa_{SC}$ | -0.144 or -0.44 | $\text{h}^{-1}$ | coupling strength from shell to core (sparse or dense) |
| $\kappa_{CC}$ | 0.5 | $\text{h}^{-1}$ | coupling strength from core to core |
| $\kappa_{SS}$ | 0.5 | $\text{h}^{-1}$ | coupling strength from shell to shell |
| $N_C$ | $13 \times 13$ | — | total number of core neurons |
| $N_S$ | $16 \times 16$ | — | total number of shell neurons |
| $n_i$ | 14, 15, 16 | — | number of connections for neuron $i$ in a subregion<br>(corners, sides or bulk of the 2D lattice) |
| $N_{CS}$ | $N_S$ | — | total number of connections from core to shell |
| $N_{SC}$ | 39 or 169 | — | total number of connections from shell to core (sparse or dense) |

Next, we investigate the after-effect. Because the intrinsic SCN period is set to 24 hours in the simulations (Table 2), no phase difference forms under T24 (Fig. 6D). In contrast, the SCN neural network develops phase differences between the core and shell regions under T22 and T26, and exhibits a prolonged after-effect in the subsequent DD when repulsive coupling from shell to core neurons is included (Fig. 6D and Fig. S4). In DD following T22 and T26, phase values are spatially homogeneous in the shell region but non-homogeneous in the core region (Fig. 6D). Since *χ* = 0.23, only a few core neurons receive repulsive coupling from shell neurons, and their phases are advanced (after T22; Fig. S4) or delayed (after T26) from the majority.

As connectivity becomes dense, a smaller magnitude of *κ*_*SC*_ causes a prolonged after-effect (Fig. S3). At *χ* = 169*/*169 = 1, the phase values within each of the subregions are spatially homogeneous during after-effect in DD (Fig. S5).

In this SCN network model, we observe crossovers of collective period in DD. Namely, at earlier time points, the collective period changes as 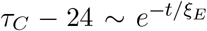, while at later time points, 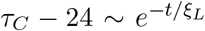 in Fig. 6E, F. These *ξ*_*E*_ and *ξ*_*L*_ are obtained by exponential fitting over 0 ∼ 10 days and 90 ∼ 100 days, respectively. With a sparse connectivity of *χ* = 0.23, the rate of change in the collective period is larger at earlier time points than at later time points, *ξ*_*E*_ *< ξ*_*L*_ (Fig. 6E). In contrast, for *χ* = 1, we observe a slower change of the collective period at earlier time points, *ξ*_*E*_ *> ξ*_*L*_ (Fig. 6F). Since SCN rhythms are reflected in mouse activity rhythms [5], we expected similar crossovers to appear in Fig. 1. However, changes in periods in DD after T-cycles could be fitted by a single exponential function, and no such crossovers were detected in these mice (Fig. 1A, B). Detecting these subtle changes may require timeseries data of the period with higher temporal resolution.

In summary, we confirmed that the combination of attractive and repulsive couplings between the core and shell regions enhances SCN entrainability to LD cycles and induces the after-effect in the SCN neural network model. Connectivity from the shell to core regions determines phase patterns in the core region and the crossover of FRP in DD.

## Discussion

Entrainment of the circadian clock to non-24 T-cycles and the resulting after-effect have been studied by using coupled phase oscillator models of the SCN. Several previous theoretical studies explained the after-effect as long transient dynamics of coupled oscillators [37, 38]. In these models, the phase difference between oscillators gradually relaxes back to its steady state value, and the FRP converges to the intrinsic period. In contrast, a recent study proposed that non-24 T-cycles alter coupling strengths between the core and shell regions of the SCN through changes in DNA methylation of genes associated with neurotransmitter receptors and ion channels [33, 34]. In that model [34], the after-effect was generated not as a transient state but as distinct frequency-locked steady states with different coupling strengths between the core and shell regions. These two mechanisms proposed in earlier studies [34, 37, 38] are not mutually exclusive; slow convergence of the FRP may occur alongside changes in the coupling strength between SCN neurons.

Although these earlier models addressed how the after-effect arises, its biological significance remained unresolved. In this study, we developed a theoretical framework based on experimentally observed phase differences between the core and shell regions of the SCN [34]. We demonstrated that repulsive coupling from shell to core regions extends the entrainment range for non-24 T-cycles. In the presence of such repulsive coupling, the shell phase “pushes” the core phase toward the light phase, bringing the SCN period closer to the LD period. This effect of repulsive coupling on entrainment may operate not only in the SCN circadian network but also in other neuronal pacemaker systems [49]. Moreover, when the magnitude of the repulsive coupling is close to that of the attractive coupling from core to shell regions, the repulsive coupling prolongs the transient dynamics of the phase difference, producing the after-effect. Repulsive coupling from the shell to core regions increases the instantaneous frequency of the core region when its phase leads the shell phase and decreases it when delayed. However, a slightly stronger attractive coupling from core to shell regions ultimately synchronizes the two SCN regions in DD. Thus, the SCN can adapt to non-24-hour T-cycles through phase difference formation and the combined action of attractive and repulsive coupling, creating a long-term memory of the experienced LD period. Beyond the after-effect, Myung et al. also used a model incorporating repulsive coupling to explain SCN adaptation to day-length changes [25]. Splitting of circadian activity rhythms in rodents under constant light condition has likewise been explained by repulsive coupling [50, 51] or by time-delayed coupling that effectively acts as a repulsive coupling [52]. Taken together, the emergence of repulsive coupling may be an adaptive mechanism of the SCN to LD cycles with large period mismatches or extended day-length.

The molecular basis underlying attractive and repulsive couplings has been partially clarified. Previous studies have shown that interactions mediated by VIP and AVP act as attractive coupling between SCN neurons [18, 21, 53]. In contrast, GABA has been proposed to mediate repulsive coupling, as GABA signaling reduces synchronization between SCN neurons [26]. Numerical simulations have also shown that tonic GABA increases phase differences between coupled neuronal oscillators [24]. Notably, treatment with a GABA receptor antagonists (gabazine) abolished correlations between SCN and T-cycle periods [34]. Assuming that GABA signaling functions as repulsive coupling, our theory predicts that blocking GABA production in the shell region would shorten the after-effect. Conversely, blocking GABA production in the core region may increase the relative influence of VIP over GABA signaling. This may result in an increase of attractive coupling from the core to shell regions *κ*_*CS*_. Hence, the phase difference between the core and shell regions generated by non-24 T-cycles should decrease according to our theory.

We showed that differences in intrinsic periods between the core and shell regions produce variations in after-effect duration depending on the T-cycle to which animals are exposed (Fig. 5). A dissection experiment demonstrated that the dorsal (shell) region of the mouse SCN has a shorter period than the ventral (core) region [43]. Based on our model, a shorter shell period relative to the core results in a longer after-effect following T26 than T22 (Fig. 5A, C). This prediction can be tested experimentally by comparing after-effect durations across T-cycles in future studies.

The SCN network model indicated that changes in FRP in DD after a non-24 T-cycle depend on the connectivity between shell and core neurons in the SCN (Fig. 6E, F). A gradually slowing rate of FRP change suggests sparse connectivity (Fig. 6E), whereas a relatively constant rate of change suggests dense connectivity (Fig. 6F). A previous theoretical and experimental study using mouse SCN slices inferred sparse connectivity from shell to core neurons [48]. If the intact SCN also maintains sparse connectivity from the shell to core regions, our model predicts that FRP changes in mice will gradually slow down during DD. This prediction could be validated experimentally using high–temporal-resolution timeseries of FRP from actograms.

We described behavioral after-effects in mice by assuming that SCN period positively correlates with behavioral period. However, earlier studies reported that the periods of *Per1/2::luc* rhythms in SCN slices from entrained animals were negatively correlated with the T-cycle period [9, 34, 36]. Specifically, SCN slices from mice entrained to T22 exhibited longer periods than those from mice entrained to T26. A recent imaging study revealed that the period of the core region is positively correlated with the T-cycle period, whereas that of the shell region is negatively correlated [34]. Moreover, the phase difference between the core and shell regions in SCN slice cultures increased over time [34]. Our current theory is based on the phase patterns of SCN slices immediately after preparation [34], which should reflect the phase distribution of the intact SCN. The process of slice preparation may alter network topology and coupling strength between neurons [34,54]. These changes could become more influential in the long-term culture experiment than the short-term one. To reproduce long-term slice dynamics, the present phase oscillator model may require additional parameter adjustments. As the three-dimensional architecture of the SCN becomes increasingly well characterized [54, 55], developing a mathematical model of the 3D SCN and simulating slice preparation [54] represent promising future directions for understanding the differences between intact and cultured SCN dynamics.

Entrainment experiments using non-24 T-cycles provide a useful paradigm for examining how the circadian clock responds to environmental cues. Because the LD period on Earth remains constant throughout a mammal ‘s lifespan, the biological significance of the after-effect has long been unclear. Our theory suggests that the after-effect arises as a consequence of enhanced circadian entrainability driven by the combined action of attractive and repulsive coupling. Interestingly, LD cycles with long day-length also induce an after-effect: mice exposed to long days exhibit shorter FRPs in subsequent DD than those exposed to short days [3]. Long-day conditions also generate complex phase patterns in the SCN that are absent under short-day conditions [11, 25, 56], suggesting that day-length information may be encoded in SCN phase patterns. Thus, phase organization within the SCN may serve as a tissue-level memory system for environmental variation, complementing molecular and cellular memory mechanisms such as DNA methylation and synaptic plasticity.

## Methods

### Data and simulation fitting

Previous experimental studies recorded mouse behavioral rhythms in DD after entrainment to T20 or T28 and reported gradual FRP changes over several months [3, 35]. We reanalyzed these FRPs to compare them with those generated by our mathematical models. Data from references [3,35] were extracted using WebPlotDigitizer ver4.7 (https://automeris.io). FRPs were averaged over individual mice and plotted on a log-linear scale from day 0 to day 41 (Fig. 1A) and to day 42 (Fig. 1B). In Fig. 1, we assumed that mouse FRPs converge to the mean of the averaged FRPs after T20 and T28. To estimate the characteristic time of the after-effect, we subtracted this mean value from daily FRP data (Fig. 1A, B) and fitted an exponential function *e*^*t/ξ*^. We used the fit and lsqcurvefit functions in the Curve Fitting Toolbox of MATLAB (The MathWorks, Natick, MA) for data and simulation fitting in Figs. 1 and 6.

### Simulations

Ordinary differential equations were numerically solved using the 4th-order Runge-Kutta (RK) method implemented through custom MATLAB scripts. The time step Δ*t* was 0.01 h for the two-oscillator model and 0.05 h for the SCN network model.

### Unit of phase difference

In several figures, we plotted the phase difference between the core and shell regions by the unit of hour (e.g. Figs. 2, 4 and 5). In Fig. 2C, we divided the phase difference *ψ*_*CS*_ by *ω*_*L*_ to fit the model to previous experimental data [34]. In Figs. 4B, 5A, 5B, S1 and S4, we divided *ψ*_*CS*_ by 2*π/*24 in both a LD cycle and DD for the continuity of phase difference at *t* = 0 (from LD to DD).

## Data availability

All data generated or analyzed during this study are included in this published article.

## Code availability

The underlying code for this study (the two-oscillator model and SCN network model) is available in Supplementary Information.

## Author contributions

Y.K., H.T. and K.U. designed research; Y.K., S.K. and K.U. performed research; H.T. and K.U. supervised research; and Y.K, H.T. and K.U. wrote the paper.

## Acknowledgments

This work was supported by JST SPRING Grant Number JPMJSP2135 to YK, JSPS KAKENHI Grant Numbers 19H04955 and 24H00863 to KU, and JSPS KAKENHI Grant Number 23K27213 to HT.

