## Supplementary Information for "Attractive and repulsive couplings between circadian pacemaker neurons promote entrainment to light-dark cycles with after-effect"

#### This PDF file includes:

Supplementary text
Figs. S1 to S5
SI References

### Supplementary Information Text

#### S1. Phase difference between the core and shell regions in an LD cycle

We derive the phase difference between core and shell regions in a light-dark (LD) condition Eq. (2) in the main text. The shell region can be entrained to an LD cycle by interacting with the core region. After entrainment, the frequency of the shell region matches that of the LD cycle

$$\frac{d\phi_L}{dt} = \omega_L \,. \quad \text{[S1]}$$

We define the phase difference between the LD cycle and the shell region $\psi_{LS} \equiv \phi_L - \phi_S$, and between the core and the shell regions $\psi_{CS} \equiv \phi_C - \phi_S$. We compute the time derivative of $\psi_{LS}$

$$\frac{d\psi_{LS}}{dt} = \omega_L - (\omega + \kappa_{CS}\sin\psi_{CS}) \,, \quad \text{[S2]}$$

where we used Eq. (S1) and the time derivative of $\phi_S$ in Eq. (1) in the main text. When the shell region is entrained to the LD cycle, the phase difference satisfies $d\psi_{LS}/dt = 0$, indicating

$$\omega_L - (\omega + \kappa_{CS}\sin\psi_{CS}) = 0 \,. \quad \text{[S3]}$$

By solving Eq. (S3), we obtain the phase difference as in Eq. (2) in the main text. Eq. (S3) has solutions only when

$$\left|\frac{\omega_L - \omega}{\kappa_{CS}}\right| \le 1, \quad \text{[S4]}$$

which determines the entrainment condition of the shell region.

#### S2. Entrainment of the core region to an LD cycle

Next, we calculate the entrainment condition of the core region to an LD cycle. Introducing the phase difference between the LD cycle and the core region $\psi_{LC} = \phi_L - \phi_C$, the differential equation of the core phase in Eq. (1) in the main text is rewritten as

$$\frac{d\phi_C}{dt} = \omega + \kappa_{SC}\sin(-\psi_{CS}) + \kappa_L \sin\psi_{LC} \,. \quad \text{[S5]}$$

Then, the time derivative of $\psi_{LC}$ is,

$$\frac{d\psi_{LC}}{dt} = \omega_L - (\omega - \kappa_{SC}\sin\psi_{CS} + \kappa_L\sin\psi_{LC}) \,. \quad \text{[S6]}$$

When the core region is entrained to the LD cycle, $d\psi_{LC}/dt = 0$. Therefore, we obtain

$$\omega_L - \omega + \kappa_{SC}\sin\psi_{CS} = \kappa_L \sin\psi_{LC} \,. \quad \text{[S7]}$$

By using Eq. (2) in the main text as $\sin\psi_{CS} = (\omega_L - \omega)/\kappa_{CS}$, we obtain

$$\frac{\omega_L - \omega}{\kappa_L}\left(1 + \frac{\kappa_{SC}}{\kappa_{CS}}\right) = \sin\psi_{LC} \,. \quad \text{[S8]}$$

Because $-1 \le \sin\psi_{LC} \le 1$, Eq. (S8) has solutions, if

$$\left|\frac{\omega_L - \omega}{\kappa_L}\left(1 + \frac{\kappa_{SC}}{\kappa_{CS}}\right)\right| \le 1 \,, \quad \text{[S9]}$$

which is the condition Eq. (4) in the main text. The phase difference $\psi_{LC}$ is

$$\psi_{LC} = \sin^{-1}\left(\frac{\omega_L - \omega}{\kappa_L}\left(1 + \frac{\kappa_{SC}}{\kappa_{CS}}\right)\right) \,. \quad \text{[S10]}$$

#### S3. Derivation of equation for phase difference between the core and shell regions in DD

In the constant dark condition (DD), we set $\kappa_L = 0$ in Eq. (1) in the main text. First, we derive the collective frequency $\Omega = (\dot{\phi}_C + \dot{\phi}_S)/2$. By substituting Eq. (1) in the main text, we obtain

$$\Omega = \frac{\dot{\phi}_C + \dot{\phi}_S}{2} = \omega + \frac{\kappa_{CS} - \kappa_{SC}}{2}\sin\psi_{CS} \,. \quad \text{[S11]}$$

Then, the time evolution of the phase difference between the core and the shell regions $\psi_{CS}$ is

$$\frac{d\psi_{CS}}{dt} = -(\kappa_{CS} + \kappa_{SC})\sin\psi_{CS} \,. \quad \text{[S12]}$$

In principle, Eq. (S12) can be solved analytically (1, 2). However, it may be difficult to understand the mechanism for the after-effect from the exact solution because of its complicated expression (see Text S5 below). Therefore, we simplify Eq. (S12) by linearization around the synchronized state $\overline{\psi}_{CS} = 0$. We assume $|\psi_{CS}| \ll 1$ and approximate $\sin\psi_{CS} \approx \psi_{CS}$ in Eqs. (S11) and (S12). Then, we obtain Eqs. (7) and (9) in the main text.

### S4. Determination of the phase shift

We consider the difference in intrinsic frequencies between the core and shell regions $\omega_C \neq \omega_S$. We introduce a phase shift $\alpha$ in Eq. (1) in the main text to modulate the steady state phase difference. The time derivatives of $\phi_C$ and $\phi_S$ in DD are

$$\frac{d\phi_C(t)}{dt} = \omega_C + \kappa_{SC}\sin(\phi_S - \phi_C - \alpha)\,,$$
$$\frac{d\phi_S(t)}{dt} = \omega_S + \kappa_{CS}\sin(\phi_C - \phi_S)\,.$$

As in the previous sections, we consider the phase difference $\psi_{CS} = \phi_C - \phi_S$. The time derivative of $\psi_{CS}$ is

$$\frac{d\psi_{CS}(t)}{dt} = \Delta\omega - \kappa_{SC}\sin(\psi_{CS} + \alpha) - \kappa_{CS}\sin\psi_{CS}\,, \tag{S13}$$

where $\Delta\omega = \omega_C - \omega_S$. By setting $d\psi_{CS}/dt = 0$, we obtain the steady state phase difference $\overline{\psi}_{CS}$. If $\alpha = 0$, the steady-state phase difference is

$$\overline{\psi}_{CS} = \sin^{-1}\left(\frac{\Delta\omega}{\kappa_{SC} + \kappa_{CS}}\right)\,. \tag{S14}$$

Hence, $\overline{\psi}_{CS} \neq 0$ for $\Delta\omega \neq 0$. However, a nonzero steady state phase difference may make it difficult to compare the results for different values of $\Delta\omega$.

Instead, we impose $\overline{\psi}_{CS} = 0$ by setting $\alpha$ to satisfy

$$\Delta\omega - \kappa_{SC}\sin\alpha = 0\,. \tag{S15}$$

Hence, the value of $\alpha$ depends on the frequency difference

$$\alpha = \sin^{-1}\left(\Delta\omega/\kappa_{SC}\right)\,. \tag{S16}$$

The linear stability analysis indicates that $\overline{\psi}_{CS} = 0$ is stable for

$$\kappa_{SC}\cos\alpha + \kappa_{CS} > 0\,. \tag{S17}$$

### S5. Analytical solution of phase difference in the presence of intrinsic frequency difference

We analytically solve the differential equation for the phase difference between the core and shell regions with a frequency mismatch

$$\frac{d\psi_{CS}}{dt} = \Delta\omega - \kappa_{SC}\sin(\psi_{CS} + \alpha) - \kappa_{CS}\sin\psi_{CS}\,, \tag{S18}$$

where $\Delta\omega = \omega_C - \omega_S \neq 0$ and $\alpha$ is defined in Eq. (S16). We separate variables and integrate both the left and right-hand sides

$$\int_{\psi_{CS}(0)}^{\psi_{CS}(t)} \frac{d\psi_{CS}}{\Delta\omega - \kappa_{SC}\sin(\psi_{CS} + \alpha) - \kappa_{CS}\sin\psi_{CS}} = \int_0^t dt\,. \tag{S19}$$

We change the variable $u = \tan(\psi_{CS}/2)$ (1). Then, we use the relations $\cos^2(\psi_{CS}/2) = (u^2+1)^{-1}$, $\cos\psi_{CS} = 2(u^2+1)^{-1} - 1$, $\sin\psi_{CS} = 2u/(u^2+1)$ and $\sin(\psi_{CS} + \alpha) = 2u(u^2+1)^{-1}\cos\alpha + \{2(u^2+1)^{-1} - 1\}\sin\alpha$,

$$\frac{1}{\Delta\omega + \kappa_{SC}\sin\alpha}\int_{u_0}^{u_1} \frac{2}{u^2 - \frac{2(\kappa_{SC}\cos\alpha + \kappa_{CS})}{\Delta\omega + \kappa_{SC}\sin\alpha}u + \frac{\Delta\omega - \kappa_{SC}\sin\alpha}{\Delta\omega + \kappa_{SC}\sin\alpha}}du = t\,, \tag{S20}$$

where $u_0 = \tan(\psi_{CS}(0)/2)$ and $u_1 = \tan(\psi_{CS}(t)/2)$. Because $\sin\alpha = \Delta\omega/\kappa_{SC}$, Eq. (S20) becomes

$$\int_{u_0}^{u_1} \frac{1}{u^2 - \frac{\kappa_{SC}\cos\alpha + \kappa_{CS}}{\Delta\omega}u}du = \Delta\omega t\,. \tag{S21}$$

After integrating the left-hand side, we obtain

$$\left[\log\left|1 - \frac{\kappa_{SC}\cos\alpha + \kappa_{CS}}{u\Delta\omega}\right|\right]_{u_0}^{u_1} = (\kappa_{SC}\cos\alpha + \kappa_{CS})\,t\,. \tag{S22}$$

$\psi_{CS}(t)$ keeps the same sign as $\psi_{CS}(0)$ and asymptotically approaches zero as $t \to \infty$ in our simulation. Hence, $u_0$ and $u_1$ are always the same sign as well. Because $\Delta\omega$ can take a negative or positive value, there are four cases:

1. $u_1 > 0, u_0 > 0$ and $\Delta\omega < 0$
2. $u_1 < 0, u_0 < 0$ and $\Delta\omega > 0$
3. $u_1 > 0, u_0 > 0$ and $\Delta\omega > 0$

4. $u_1 < 0, u_0 < 0$ and $\Delta\omega < 0$

Based on our simulation parameters, $1 < |(\kappa_{SC}\cos\alpha + \kappa_{CS})/(u_1\Delta\omega)|$ and $1 < |(\kappa_{SC}\cos\alpha + \kappa_{CS})/(u_0\Delta\omega)|$ are satisfied. Therefore, for all these four cases, Eq. (S22) becomes

$$\log\left(-1 + \frac{\kappa_{SC}\cos\alpha + \kappa_{CS}}{u_1\Delta\omega}\right)\left(-1 + \frac{\kappa_{SC}\cos\alpha + \kappa_{CS}}{u_0\Delta\omega}\right)^{-1} = (\kappa_{SC}\cos\alpha + \kappa_{CS})\,t\,. \qquad \text{[S23]}$$

We solve this equation for $u_1$

$$u_1 = -\frac{u_0}{u_0 + (\frac{\kappa_{SC}\cos\alpha+\kappa_{CS}}{\Delta\omega} - u_0)e^{(\kappa_{SC}\cos\alpha+\kappa_{CS})t}}\frac{\kappa_{SC}\cos\alpha + \kappa_{CS}}{\Delta\omega}\,. \qquad \text{[S24]}$$

Thus, $\psi_{CS}(t)$ is

$$\psi_{CS}(t) = 2\tan^{-1}\left(\frac{(\kappa_{SC}\cos\alpha + \kappa_{CS})\tan\frac{\psi_{CS}(0)}{2}}{\Delta\omega\tan\frac{\psi_{CS}(0)}{2} + (\kappa_{SC}\cos\alpha + \kappa_{CS} - \Delta\omega\tan\frac{\psi_{CS}(0)}{2})e^{(\kappa_{SC}\cos\alpha+\kappa_{CS})t}}\right)\,. \qquad \text{[S25]}$$

To understand the relation between a prolonged after-effect and $\Delta\omega$, we consider the denominator of the argument of the arctangent in Eq. (S25), because the numerator does not include time $t$

$$\Delta\omega\tan\frac{\psi_{CS}(0)}{2} + (\kappa_{SC}\cos\alpha + \kappa_{CS} - \Delta\omega\tan\frac{\psi_{CS}(0)}{2})e^{(\kappa_{SC}\cos\alpha+\kappa_{CS})t}\,. \qquad \text{[S26]}$$

If $\Delta\omega\tan(\psi_{CS}(0)/2) < 0$, the relation Eq. (S17) leads to

$$|\Delta\omega\tan(\psi_{CS}(0)/2)| < \kappa_{SC}\cos\alpha + \kappa_{CS} - \Delta\omega\tan(\psi_{CS}(0)/2)\,, \qquad \text{[S27]}$$

indicating that the second term of Eq. (S26) is already dominant at $t = 0$. Hence, $\psi_{CS}(t)$ immediately decreases, suggesting a shorter after-effect.

In contrast, if $\Delta\omega\tan(\psi_{CS}(0)/2) > 0$, the second term of Eq. (S26) could be much smaller than the first term at $t = 0$,

$$\Delta\omega\tan(\psi_{CS}(0)/2) \gg \kappa_{SC}\cos\alpha + \kappa_{CS} - \Delta\omega\tan(\psi_{CS}(0)/2) > 0\,. \qquad \text{[S28]}$$

In this case, change in $\psi_{CS}(t)$ would be very small until roughly

$$t_c = \frac{1}{\kappa_{SC}\cos\alpha + \kappa_{CS}}\log\left(\frac{\Delta\omega\tan(\psi_{CS}(0)/2)}{\kappa_{SC}\cos\alpha + \kappa_{CS} - \Delta\omega\tan(\psi_{CS}(0)/2)}\right)\,. \qquad \text{[S29]}$$

Thus, $\Delta\omega\tan(\psi_{CS}(0)/2) > 0$ is a condition for a prolonged after-effect. When $\Delta\omega < 0$, we observed a longer after-effect for T26 which resulted in $\psi_{CS}(0) < 0$ (Fig. 5A). When $\Delta\omega > 0$, $\psi_{CS}(0) > 0$ satisfies this condition and we observed a longer after-effect for T22 in Fig. 5B in the main text.

### S6. SCN network model

We construct a SCN network model to understand the phase dynamics in individual neurons consisting of the core and shell regions. We locate SCN core neurons ($1 \le i \le N_C$) and shell neurons ($N_C < i \le N_C + N_S = N$) in square lattices (Fig. 6A). Within the core and shell regions, there are short-range couplings by diffusive neurotransmitters, such as VIP and AVP, and long-range coupling by extended synaptic connections (3–7). To describe these couplings, we use an adjacency matrix $a_{ij}$ (8). If there is a coupling from neuron $j$ to neuron $i$, $a_{ij} = 1$, otherwise $a_{ij} = 0$. We use undirected graphs for neuronal coupling within core and shell regions (i.e., if $a_{ij} = 1$, then $a_{ji} = 1$). Neurons in the bulk of the core or shell region are coupled to the four nearest neighbors, and those at the boundary are coupled to their two (corner) or three (sides of the square) nearest neighbors. In addition, each neuron is connected to 12 randomly chosen neurons within the same regions.

To describe the interactions between core and shell neurons, we use directed graphs. All shell neurons receive a single directed connection from a randomly chosen core neuron. The total number of connections from core neurons to shell neurons is $N_S$. If $N_S > N_C$, $N_S - N_C$ core neurons send links to two different shell neurons and $N_C - (N_S - N_C)$ core neurons send a single link to a shell neuron. In contrast, only $N_{SC}$ core neurons receive a directed connection from a randomly chosen shell neuron. If $N_{SC} < N_C$, $(N_C - N_{SC})$ core neurons are not coupled to any shell neuron. See the subsection below for the algorithms of SCN network construction.

We describe the time evolution of the phase of individual core neurons $\phi_i$ for $1 \le i \le N_C$, the phase of individual shell neurons $\phi_i$ for $N_C < i \le N_C + N_S = N$ and the phase of the LD cycle $\phi_L$ as

$$\frac{d\phi_i}{dt} = \frac{2\pi}{24} + \frac{\kappa_{CC}}{n_i}\sum_{j=1}^{N_C} a_{ij}\sin(\phi_j - \phi_i) + \kappa_{SC}\sum_{j=N_C+1}^{N_C+N_S} a_{ij}\sin(\phi_j - \phi_i) + \kappa_L \sin(\phi_L - \phi_i)\ (\text{for } 1 \le i \le N_C), \quad \text{[S30]}$$

$$\frac{d\phi_i}{dt} = \frac{2\pi}{24} + \kappa_{CS}\sum_{j=1}^{N_C} a_{ij}\sin(\phi_j - \phi_i) + \frac{\kappa_{SS}}{n_i}\sum_{j=N_C+1}^{N_C+N_S} a_{ij}\sin(\phi_j - \phi_i) \quad \text{[S31]}$$

$$(\text{for } N_C < i),$$

$$\frac{d\phi_L}{dt} = \omega_L, \quad \text{[S32]}$$

where $n_i$ is the number of core or shell neurons coupled to neuron $i$, $n_i = \sum_{j=1}^{N_C} a_{ij}$ for $1 \le i \le N_C$, and $n_i = \sum_{j=N_C+1}^{N_C+N_S} a_{ij}$ for $N_C < i$. The first terms in Eqs. (S30) and (S31) represent the intrinsic frequency of core and shell neurons. In Eq. (S30), the second and third terms are coupling from core neurons and a shell neuron, respectively, to core neuron $i$. The fourth term in Eq. (S30) is coupling to the light phase. In Eq. (S31), the second and third terms are coupling from a core neuron and from shell neurons, respectively, to shell neuron $i$. We define coupling strengths from core to core $\kappa_{CC}$, shell to core $\kappa_{SC}$, light to core $\kappa_L$, core to shell $\kappa_{CS}$ and shell to shell $\kappa_{SS}$ in Eqs. (S30) and (S31). For simplicity, we assume the same coupling strength $\kappa_{CC} = \kappa_{SS}$. $\omega_L$ in Eq.(S32) is the frequency of the LD cycle.

To obtain the average SCN phase $\overline{\phi}$, we first compute Kuramoto order parameter (9)

$$re^{i\overline{\phi}} = r(\cos\overline{\phi} + i\sin\overline{\phi}) = \frac{1}{N}\sum_{j=1}^{N} e^{i\phi_j}, \quad \text{[S33]}$$

where

$$r\cos\overline{\phi} = \frac{1}{N}\sum_{j=1}^{N}\cos\phi_j\ , \quad \text{[S34]}$$

$$r\sin\overline{\phi} = \frac{1}{N}\sum_{j=1}^{N}\sin\phi_j\ . \quad \text{[S35]}$$

Then, we calculate $\overline{\phi}$ from the relation

$$\tan\overline{\phi} = \frac{\sin\overline{\phi}}{\cos\overline{\phi}}. \quad \text{[S36]}$$

The phase difference between neuron $i$ and the average SCN phase is defined as

$$\psi_i \equiv \phi_i - \overline{\phi}. \quad \text{[S37]}$$

Similarly, we define average core and shell phases $r_C e^{i\overline{\phi_C}} = N_C^{-1}\sum_{j=1}^{N_C} e^{i\phi_j}$ and $r_S e^{i\overline{\phi_S}} = N_S^{-1}\sum_{j=N_C+1}^{N} e^{i\phi_j}$, respectively. We then calculate the average phase difference between core and shell $\overline{\psi_{CS}} = \overline{\phi_C} - \overline{\phi_S}$ in Fig. 6D in the main text and Fig. S4. In addition, the period in SCN network model at each time point is calculated by $2\pi\{(\dot{\overline{\phi_C}} + \dot{\overline{\phi_S}})/2\}^{-1}$ in Fig. 6E and F in the main text.

**Algorithms of generating SCN networks.** Here we describe how we make the adjacency matrix $a_{ij}$ $(N \times N)$ that determines the SCN network topology. First, we set all the entries of the matrix to zero. In this matrix, the left upper $N_C \times N_C$ region represents coupling from a core neuron to another core neuron. We assign $(X, Y)$ coordinates to these core neurons by $X = \text{ceil}(i/\sqrt{N_C})$, $Y = \text{mod}(i-1, \sqrt{N_C}) + 1$ where ceil is the ceil function rounding up to the nearest integer number. We calculate the distance for a pair of core neurons $i$ and $j$, $r = \sqrt{(X_i - X_j)^2 + (Y_i - Y_j)^2}$, and if $r = 1$, $a_{ij} = 1$ and $a_{ji} = 1$ to make nearest neighbor coupling. Next, for neuron $i$, we randomly choose $n_R = 12$ core neurons $j_1, .., j_{n_R}$ and set $a_{ij_k} = 1$ and $a_{j_k i} = 1$ if $r_{ij_k} > 1$. If some core neurons have connections larger or less than $n_R$, we discard the adjacency matrix $a_{ij}$ and go back to the previous step. We do the same procedures for the entry for the shell neurons (right bottom $(N - N_C) \times (N - N_C)$ region of the matrix).

Next, we determine the $a_{ij}$ of the left bottom part $(N - N_C) \times N_C$ region representing coupling from a core neuron to a shell neuron. We first make $N_C$ random pairs of core and shell neurons without overlapping. These are directed connections, so $a_{ij} = 1$ for core neuron $j$ and shell neuron $i$. Because $N - N_C = N_S$ is larger than $N_C$ in our simulation, we further make core-shell pairs for the remaining $N_S - N_C$ shell neurons, by using overlapping $N_S - N_C$ core neurons. This means that $N_S - N_C$ core neurons send phase information to two shell neurons ($\sum_{i=N_C+1}^{N} a_{ij} = 2$), while $2N_C - N_S$ core neurons do to one shell neuron ($\sum_{i=N_C+1}^{N} a_{ij} = 1$).

We do the same procedures for the coupling from $N_{SC}$ shell neurons to $N_{SC}$ core neurons to determine the right upper $N_C \times (N - N_C)$ region of the adjacency matrix. In our simulation, $N_S$ is larger than $N_{SC}$. This means that each of $N_{SC}$ shell neurons sends phase information to a core neuron ($\sum_{i=1}^{Nc} a_{ij} = 1$), whereas remaining $N_S - N_{SC}$ shell neurons do not ($\sum_{i=1}^{Nc} a_{ij} = 0$).

### S7. Relation between critical coupling strength and connectivity

In this section, we derive the dependency of after-effect on the connectivity $\chi \equiv N_{SC}/N_C$ in the SCN network model. Because each neuron has 12 randomly selected connections together with nearest-neighbor connections within the core and shell regions, we expect the validity of a mean-field approximation within these regions in Eqs. (S30) and (S31),

$$\frac{d\phi_i}{dt} = \frac{2\pi}{24} + \frac{\kappa_{CC}}{N_C} \sum_{j=1}^{N_C} \sin(\phi_j - \phi_i) + \kappa_{SC} \sum_{j=N_C+1}^{N_C+N_S} a_{ij} \sin(\phi_j - \phi_i) \quad \text{(for } 1 \le i \le N_C\text{)}, \tag{S38}$$

$$\frac{d\phi_i}{dt} = \frac{2\pi}{24} + \kappa_{CS} \sum_{j=1}^{N_C} a_{ij} \sin(\phi_j - \phi_i) + \frac{\kappa_{SS}}{N_S} \sum_{j=N_C+1}^{N_C+N_S} \sin(\phi_j - \phi_i) \quad \text{(for } N_C < i \le N_C + N_S\text{)}. \tag{S39}$$

Next, based on the presence of the directed connection from shell to core, we sort neuron index $i$ for core neurons and divide them into two groups. A core neuron with index $i$ ($1 \le i \le N_{SC}$) receives a directed connection from a shell neuron. The remaining core neurons ($N_{SC} < i \le N_C$) do not receive a directed connection. After this sort, we separate summation for core neurons in Eq. (S38) into these two groups, and approximate the differential equation of a core neuron $i$ ($1 \le i \le N_{SC}$) by linearization

$$\begin{aligned} \frac{d\phi_i}{dt} &= \frac{2\pi}{24} + \frac{\kappa_{CC}}{N_C} \sum_{j=1}^{N_C} \sin(\phi_j - \phi_i) + \kappa_{SC} \sum_{j=N_C+1}^{N_C+N_S} a_{ij} \sin(\phi_j - \phi_i) \\ &\approx \frac{2\pi}{24} + \frac{\kappa_{CC}}{N_C} \left\{ \sum_{j=1}^{N_{SC}} (\phi_j - \phi_i) + \sum_{j=N_{SC}+1}^{N_C} (\phi_j - \phi_i) \right\} + \kappa_{SC} \sum_{j=N_C+1}^{N_C+N_S} a_{ij} (\phi_j - \phi_i) \, . \end{aligned} \tag{S40}$$

We do the same calculations for the remaining core neurons ($N_{SC} < i \le N_C$) and shell neurons ($N_C < i \le N_C + N_S$). We then change the notation of $\phi_i$ for the neuron $i$ ($1 \le i \le N_{SC}$) in the core region into $C_i$, which is connected to a shell neuron. The remaining neurons with the index $N_{SC} < i \le N_C$ are those unconnected to shell, and we denote each of them as $U_i$. We also change the notation of $\phi_i$ for the shell neuron $i$ into $S_i$. We obtain

$$\frac{dC_i}{dt} = \frac{2\pi}{24} + \frac{\kappa_{CC}}{N_C} \left\{ \sum_{j=1}^{N_{SC}} (C_j - C_i) + \sum_{j=N_{SC}+1}^{N_C} (U_j - C_i) \right\} + \kappa_{SC} \sum_{j=N_C+1}^{N_C+N_S} a_{ij} (S_j - C_i) \, , \tag{S41}$$

$$\frac{dU_i}{dt} = \frac{2\pi}{24} + \frac{\kappa_{CC}}{N_C} \left\{ \sum_{j=1}^{N_{SC}} (C_j - U_i) + \sum_{j=N_{SC}+1}^{N_C} (U_j - U_i) \right\} , \tag{S42}$$

$$\frac{dS_i}{dt} = \frac{2\pi}{24} + \kappa_{CS} \left\{ \sum_{j=1}^{N_{SC}} a_{ij} (C_j - S_i) + \sum_{j=N_{SC}+1}^{N_C} a_{ij} (U_j - S_i) \right\} + \frac{\kappa_{SS}}{N_S} \sum_{j=N_C+1}^{N_C+N_S} (S_j - S_i) \, . \tag{S43}$$

Also, we define average phase $\overline{C} \equiv N_{SC}^{-1} \sum_{j=1}^{N_{SC}} C_j$, $\overline{U} \equiv (N_C - N_{SC})^{-1} \sum_{j=N_{SC}+1}^{N_C} U_j$ and $\overline{S} \equiv N_S^{-1} \sum_{j=N_C+1}^{N} S_j$. Substituting these average phases into Eqs. (S41)-(S43), we obtain

$$\frac{dC_i}{dt} = \frac{2\pi}{24} + \frac{\kappa_{CC}}{N_C} \left\{ N_{SC}\overline{C} + (N_C - N_{SC})\overline{U} - N_C C_i \right\} + \kappa_{SC} \sum_{j=N_C+1}^{N_C+N_S} a_{ij} S_j - \kappa_{SC} C_i \, , \tag{S44}$$

$$\frac{dU_i}{dt} = \frac{2\pi}{24} + \frac{\kappa_{CC}}{N_C} \left\{ N_{SC}\overline{C} + (N_C - N_{SC})\overline{U} - N_{SC} U_i - (N_C - N_{SC}) U_i \right\} , \tag{S45}$$

$$\frac{dS_i}{dt} = \frac{2\pi}{24} + \kappa_{CS} \left\{ \sum_{j=1}^{N_{SC}} a_{ij} C_j + \sum_{j=N_{SC}+1}^{N_C} a_{ij} U_j - S_i \right\} + \kappa_{SS} \left( \overline{S} - S_i \right) \, . \tag{S46}$$

In Eq. (S44), we approximate $S_j \approx \overline{S}$ because the phase in shell region is almost spatially uniform in simulation (Fig. 6D),

$$\frac{dC_i}{dt} = \frac{2\pi}{24} + \frac{\kappa_{CC}}{N_C} \left\{ N_{SC}\overline{C} + (N_C - N_{SC})\overline{U} - N_C C_i \right\} + \kappa_{SC} (\overline{S} - C_i) \, . \tag{S47}$$

In Eq. (S46), we approximate $a_{ij}$ by a uniform probability $1/N_C$ for having a connection with core neuron $j$

$$\sum_{j=1}^{N_{SC}} a_{ij}C_j \approx \sum_{j=1}^{N_{SC}} \frac{1}{N_C}C_j = \frac{N_{SC}}{N_C}\overline{C}\,, \tag{S48}$$

$$\sum_{j=N_{SC}+1}^{N_C} a_{ij}U_j \approx \sum_{j=N_{SC}+1}^{N_C} \frac{1}{N_C}U_j = \frac{N_C - N_{SC}}{N_C}\overline{U}\,. \tag{S49}$$

By substituting Eqs. (S48) and (S49) into Eq. (S46), we approximate

$$\frac{dS_i}{dt} = \frac{2\pi}{24} + \kappa_{CS}\left(\frac{N_{SC}}{N_C}\overline{C} + \frac{N_C - N_{SC}}{N_C}\overline{U} - S_i\right) + \kappa_{SS}\left(\overline{S} - S_i\right)\,. \tag{S50}$$

After taking the summation for the index $i$ in Eqs. (S45), (S47) and (S50), and dividing them by $N_C - N_{SC}$, $N_{SC}$ and $N_S$, respectively, we obtain the differential equations of these three average phases

$$\frac{d\overline{C}}{dt} = \frac{2\pi}{24} + \frac{\kappa_{CC}}{N_C}(N_C - N_{SC})(\overline{U} - \overline{C}) + \kappa_{SC}(\overline{S} - \overline{C})\,, \tag{S51}$$

$$\frac{d\overline{U}}{dt} = \frac{2\pi}{24} + \frac{\kappa_{CC}}{N_C}(\overline{C} - \overline{U})\,, \tag{S52}$$

$$\frac{d\overline{S}}{dt} = \frac{2\pi}{24} + \kappa_{CS}\left(\frac{N_{SC}}{N_C}\overline{C} + \frac{N_C - N_{SC}}{N_C}\overline{U} - \overline{S}\right)\,. \tag{S53}$$

Next, we define average phase differences $\psi = \overline{U} - \overline{C}$ and $\theta = \overline{S} - \overline{C}$. We then describe the time evolution $\dot{\psi} = \dot{\overline{U}} - \dot{\overline{C}}$ and $\dot{\theta} = \dot{\overline{S}} - \dot{\overline{C}}$ as

$$\begin{pmatrix}\dot{\psi}\\ \dot{\theta}\end{pmatrix} = \begin{pmatrix} -\kappa_{CC} & -\kappa_{SC} \\ \frac{N_C - N_{SC}}{N_C}(\kappa_{CS} - \kappa_{CC}) & -(\kappa_{CS} + \kappa_{SC})\end{pmatrix}\begin{pmatrix}\psi\\ \theta\end{pmatrix}\,. \tag{S54}$$

The eigenvalues $\lambda_1, \lambda_2$ of the matrix in Eq. (S54) are

$$\lambda_{1,2} = \frac{-(\kappa_{CC} + \kappa_{CS} + \kappa_{SC}) \pm \sqrt{(\kappa_{CC} + \kappa_{CS} + \kappa_{SC})^2 - 4\{\frac{N_C - N_{SC}}{N_C}\kappa_{SC}(\kappa_{SC} - \kappa_{CC}) + \kappa_{CC}(\kappa_{SC} + \kappa_{CS})\}}}{2}.$$

Synchronized steady state $\overline{\psi} = \overline{\theta} = 0$ are stable, if $\lambda_{1,2} < 0$. In addition, the larger eigenvalue

$$\lambda_1 = \frac{-(\kappa_{CC} + \kappa_{CS} + \kappa_{SC}) + \sqrt{(\kappa_{CC} + \kappa_{CS} + \kappa_{SC})^2 - 4\{\frac{N_C - N_{SC}}{N_C}\kappa_{SC}(\kappa_{SC} - \kappa_{CC}) + \kappa_{CC}(\kappa_{SC} + \kappa_{CS})\}}}{2}\,, \tag{S55}$$

determines the speed for the phase differences to disappear. Hence, the duration of after-effect becomes longer as $\lambda_1$ approaches zero. We assume $\lambda_1 = 0$ to be the lower bound of the decay rate of the phase differences. By setting $\lambda_1 = 0$ in Eq. (S55) and solving the equation with respect to $\kappa_{SC}$, we obtain Eq. (17) in the main text

$$\kappa_{SC} = \frac{\kappa_{CC}\kappa_{CS}}{\chi(\kappa_{CS} - \kappa_{CC}) - \kappa_{CS}}\,, \tag{S56}$$

where $\chi = N_{SC}/N_C$ is the connectivity defined in Eq. (16) in the main text.

### S8. Linear stability analysis of the SCN network

In simulations, neurons in the SCN are first entrained to an LD cycle and then released in DD. To find a condition for a prolonged after-effect, we performed a linear stability analysis of the synchronized state in DD in the two-oscillator model (Text S1). The phase difference between the core and the shell regions in DD decreases at the rate determined by the eigenvalue $\lambda < 0$ of the linearized system. In the two-oscillator model, the condition for a prolonged after-effect $|\lambda| \ll 1$ is $\lambda = -(\kappa_{SC} + \kappa_{CS}) < 0$ and $|\kappa_{SC}|/\kappa_{CS} \approx 1$.

We also perform a linear stability analysis of the synchronized state in the SCN network model to explore parameter sets for a prolonged after-effect. We consider the phase difference between neuron $i$ and core neuron 1, $\psi_i = \phi_i - \phi_1$ without loss of generality. The time evolution of the small phase difference between core neuron $i$ $(1 \le i \le N_C)$ and core neuron 1 $(|\psi_i| \ll 1)$ can be approximated by using Eq. (S30)

$$\begin{aligned}\frac{d\psi_i}{dt} &= \frac{d\phi_i}{dt} - \frac{d\phi_1}{dt}\\
&= \frac{\kappa_{CC}}{n_i}\sum_{j=1}^{N_C} a_{ij}\sin(\psi_j - \psi_i) + \kappa_{SC}\sum_{j=N_C+1}^{N_C+N_S} a_{ij}\sin(\psi_j - \psi_i) - \left(\frac{\kappa_{CC}}{n_1}\sum_{j=1}^{N_C} a_{1j}\sin\psi_j + \kappa_{SC}\sum_{j=N_C+1}^{N_C+N_S} a_{1j}\sin\psi_j\right)\\
&\approx \frac{\kappa_{CC}}{n_i}\sum_{j=1}^{N_C} a_{ij}(\psi_j - \psi_i) + \kappa_{SC}\sum_{j=N_C+1}^{N_C+N_S} a_{ij}(\psi_j - \psi_i) - \left(\frac{\kappa_{CC}}{n_1}\sum_{j=1}^{N_C} a_{1j}\psi_j + \kappa_{SC}\sum_{j=N_C+1}^{N_C+N_S} a_{1j}\psi_j\right)\,.\end{aligned} \tag{S57}$$

Similarly, the time evolution of the phase difference between shell neuron $i$ ($N_C + 1 \le i \le N_C + N_S$) and core neuron 1 is approximated by using Eqs. (S30) and (S31) as

$$\begin{aligned}\frac{d\psi_i}{dt} &= \frac{d\phi_i}{dt} - \frac{d\phi_1}{dt} \\ &= \kappa_{CS}\sum_{j=1}^{N_C} a_{ij}\sin(\psi_j - \psi_i) + \frac{\kappa_{SS}}{n_i}\sum_{j=N_C+1}^{N_C+N_S} a_{ij}\sin(\psi_j - \psi_i) - \left(\frac{\kappa_{CC}}{n_1}\sum_{j=1}^{N_C} a_{1j}\sin\psi_j + \kappa_{SC}\sum_{j=N_C+1}^{N_C+N_S} a_{1j}\sin\psi_j\right) \\ &\approx \kappa_{CS}\sum_{j=1}^{N_C} a_{ij}(\psi_j - \psi_i) + \frac{\kappa_{SS}}{n_i}\sum_{j=N_C+1}^{N_C+N_S} a_{ij}(\psi_j - \psi_i) - \left(\frac{\kappa_{CC}}{n_1}\sum_{j=1}^{N_C} a_{1j}\psi_j + \kappa_{SC}\sum_{j=N_C+1}^{N_C+N_S} a_{1j}\psi_j\right).\end{aligned} \quad [S58]$$

Eqs. (S57) and (S58) can also be described by using a linearized matrix and phase difference vector $\mathbf{\Psi} = (\psi_1, ..., \psi_N)^{\mathrm{T}}$

$$\frac{d\mathbf{\Psi}}{dt} = \mathbb{B}\mathbf{\Psi}. \quad [S59]$$

$\mathbb{B}$ can be described as

$$\mathbb{B} = \mathbb{V} - \mathbb{W}, \quad [S60]$$

with

$$\mathbb{V} = \left(\begin{array}{cccc|cccc} v_{1,1} & \cdots & \cdots & v_{1,N_C} & v_{1,N_C+1} & \cdots & \cdots & v_{1,N_C+N_S} \\ \vdots & \ddots & \cdots & \vdots & \vdots & \ddots & \cdots & \vdots \\ \vdots & \cdots & \ddots & \vdots & \vdots & \cdots & \ddots & \vdots \\ v_{N_C,1} & \cdots & \cdots & v_{N_C,N_C} & v_{N_C,N_C+1} & \cdots & \cdots & v_{N_C,N_C+N_S} \\ \hline v_{N_C+1,1} & \cdots & \cdots & v_{N_C+1,N_C} & v_{N_C+1,N_C+1} & \cdots & \cdots & v_{N_C+1,N_C+N_S} \\ \vdots & \ddots & \cdots & \vdots & \vdots & \ddots & \cdots & \vdots \\ \vdots & \cdots & \ddots & \vdots & \vdots & \cdots & \ddots & \vdots \\ v_{N_C+N_S,1} & \cdots & \cdots & v_{N_C+N_S,N_C} & v_{N_C+N_S,N_C+1} & \cdots & \cdots & v_{N_C+N_S,N_C+N_S} \end{array}\right),$$

and

$$\mathbb{W} = \begin{pmatrix} \mathbf{w}_1, & \mathbf{w}_2, & \cdots & \mathbf{w}_{N_C}, & \mathbf{w}_{N_C+1}, & \cdots & \mathbf{w}_{N_C+N_S} \end{pmatrix}.$$

Each component of $\mathbb{V} = v_{ij}$ is

$$v_{ij} = \begin{cases} \frac{\kappa_{CC}}{n_i}a_{ij} & (i < N_C,\, j \le N_C) \\ \kappa_{SC}a_{ij} & (i < N_C,\, j > N_C) \\ -(\kappa_{CC} + \kappa_{SC}\sum_{j=N_C+1}^{N_C+N_S} a_{ij}) & (i \le N_C,\, j = i) \\ \kappa_{CS}a_{ij} & (i > N_C,\, j \le N_C) \\ \frac{\kappa_{SS}}{n_i}a_{ij} & (i > N_C,\, j > N_C) \\ -(\kappa_{SS} + \kappa_{CS}a_{ij}) & (i > N_C,\, j = i). \end{cases}$$

Each vector component of $\mathbb{W}$, $\mathbf{w}_j = w_{kj}$, is

$$w_{kj} = \begin{cases} -(\kappa_{CC} + \kappa_{SC}a_{11}) & (j = 1) \\ \frac{\kappa_{CC}}{n_1}a_{1j} & (1 < j \le N_C) \\ \kappa_{SC}a_{1j} & (N_C < j). \end{cases}$$

The linearized matrix $\mathbb{B}$ has $N$ eigenvalues and one of them is zero. If there is a positive eigenvalue, the synchronized state is unstable. If all the $N - 1$ eigenvalues are negative, the synchronized state is stable. In this case, we hypothesize that the second maximum eigenvalue $\lambda$ approximates the decay rate of average phase difference $\overline{\phi_C} - \overline{\phi_S}$ between the core and shell regions, and therefore the decay rate of the after-effect.

Because the condition for long after-effect is $|\lambda| \ll 1$, we seek the value of $\kappa_{SC}$ that achieves $\lambda \to -0$. To do this, we detect the zero crossing point of $\lambda$ by changing the value of $\kappa_{SC}$. We then numerically find the $\kappa_{SC}$ value for $\lambda = 0$ by linear regression analysis. In Fig. S3, we plot $|\kappa_{SC}|/\kappa_{CS}$ values obtained by this method as a function of connectivity $\chi$ (red triangles).

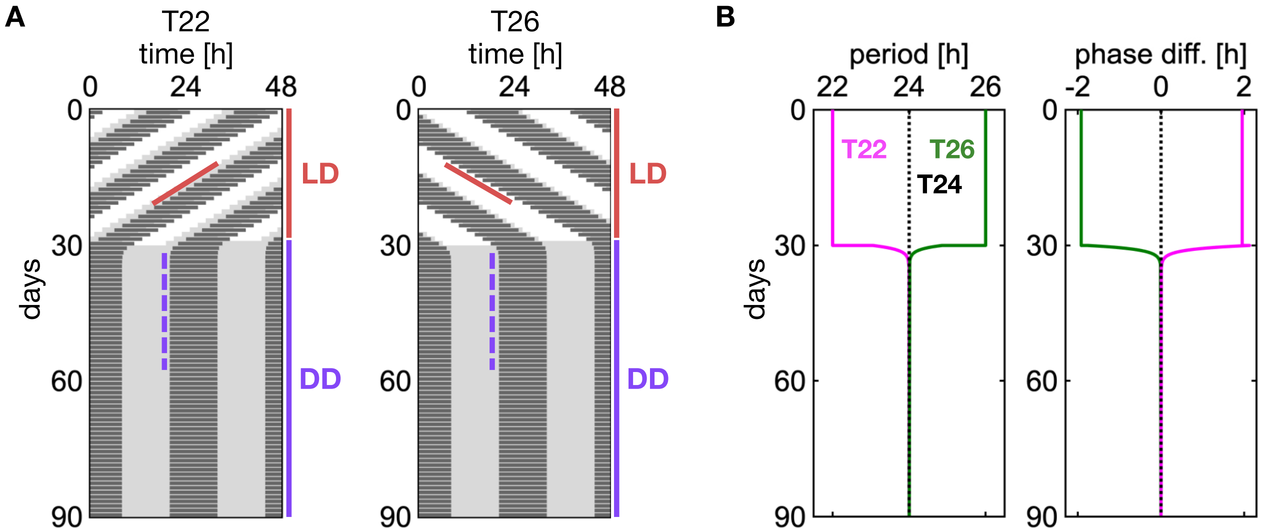


**Fig. S1.** After-effect by the combinations of attractive couplings between the core and shell regions ($\kappa_{SC}/\kappa_{CS} \approx 0.98$). (A) Simulated double-plot actogram in non-24 T-cycles (T22: left, T26: right) and subsequent DD. Black bars indicate shell phase $\phi_S$ in the interval between $\pi$ and $2\pi$, which we define as an active phase of an animal. Solid red and dotted purple slopes indicate activity onsets defined as $\phi_S = \pi$. (B) Timeseries of (left) collective period of core and shell $2\pi/\Omega$ and (right) their phase difference $\psi_{CS}$. $\Omega$ is the collective frequency $\Omega = (\dot{\phi}_C + \dot{\phi}_S)/2$. Simulation results in LD cycle from day 0 to 30 (red vertical line in (A)) and in DD from day 30 to 90 (purple vertical line in (A)). To represent SCN phase difference $\psi_{CS}$ by a unit of time (hours), we divide $\psi_{CS}$ by $2\pi/24$ in both a LD cycle and DD for continuity of phase difference at the transition at $t = 30$ (from LD to DD; Methods).

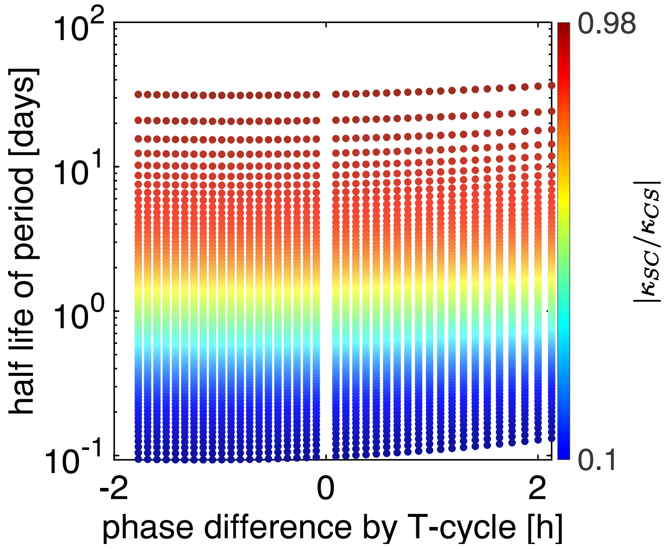


**Fig. S2.** The half-life of free running period in DD is independent of phase difference between the core and shell regions generated by a non-24 T-cycle. Colored circles indicate results for different values of $|\kappa_{SC}/\kappa_{CS}|$. We plot the steady state phase difference in non-24 T-cycles in the horizontal axis. The horizontal axis is shown on a linear scale and the vertical axis on a logarithmic scale.

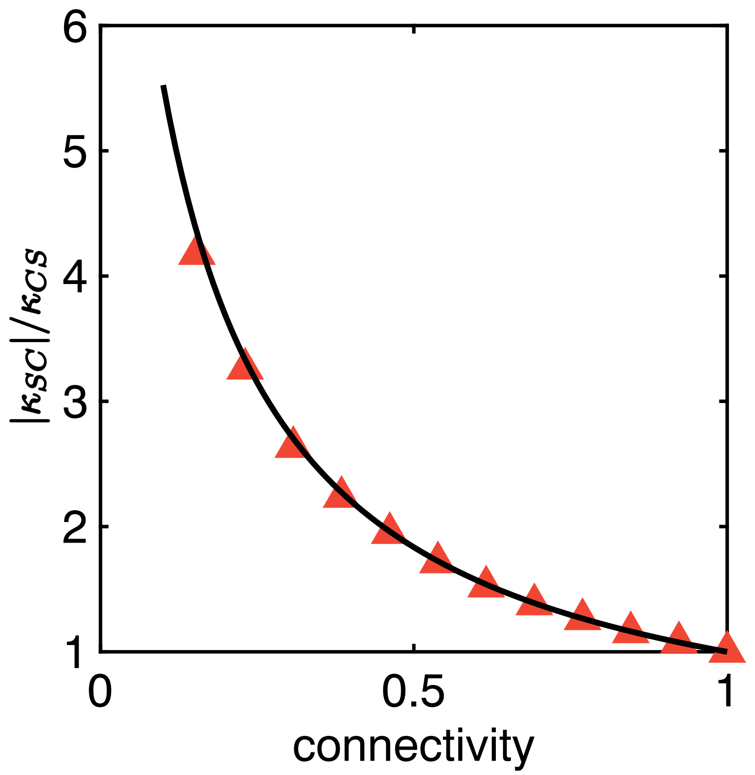


**Fig. S3.** Relationship between the connectivity $\chi$ and the critical value of $|\kappa_{SC}|/\kappa_{CS}$ where the synchronized state $\overline{\psi} = 0$ loses linear stability in the SCN network model. The black line is the analytical result Eq. (17) in the main text obtained by a mean-filed approximation. Red triangles are critical values obtained by numerical linear stability analysis of the original SCN network model. See Supplementary Information Text S7 and S8 for the details of these analyses.

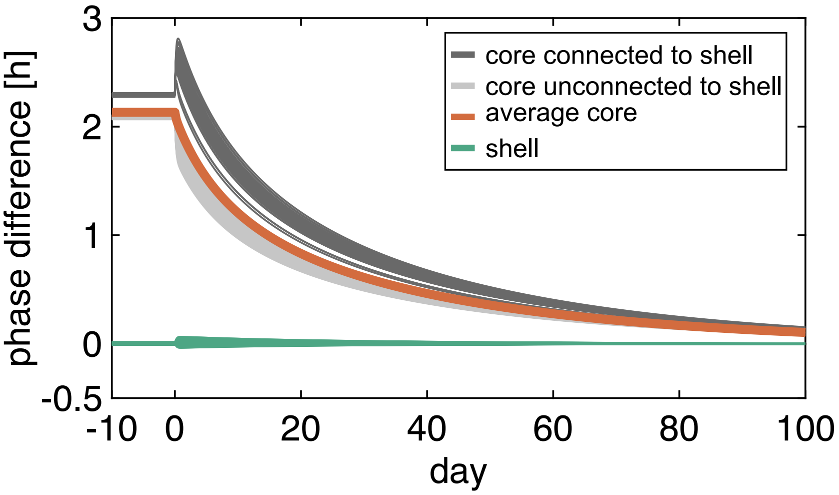


**Fig. S4.** Time-series of phase differences between individual neurons and the average of shell neurons in the SCN network model. We separate the core neurons into two groups; core neurons that receive a directed coupling from a shell neuron (dark gray), and those that do not receive a coupling (light gray). The SCN network is entrained to T22 and released in DD at $t = 0$.

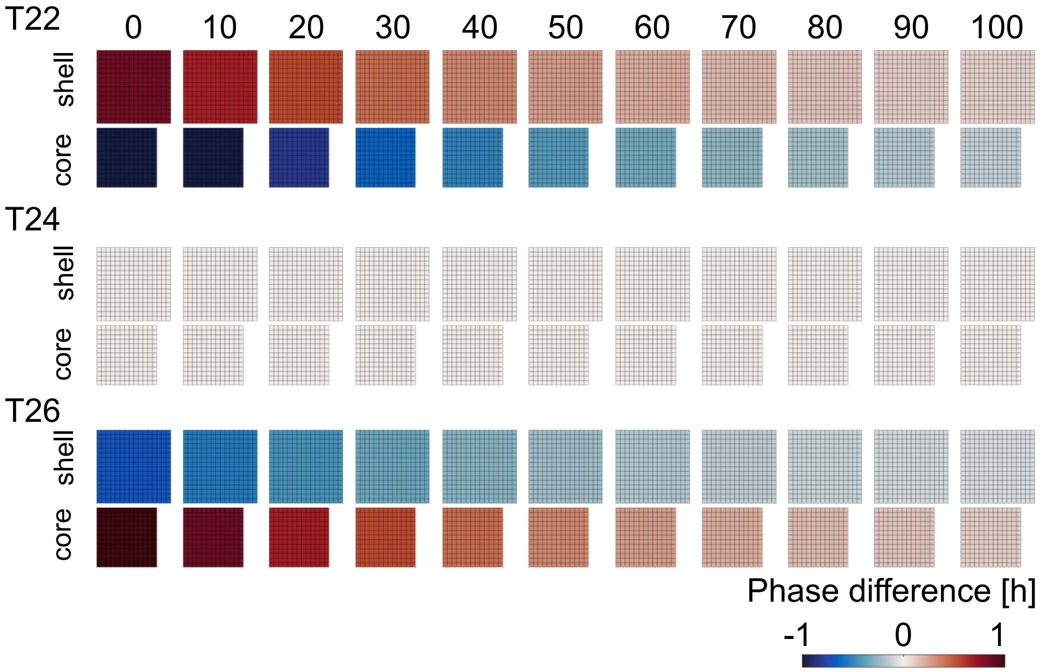


**Fig. S5.** After-effect in the SCN network model with the densest connectivity $\chi = 1$. Snapshots of phase difference between individual neurons $\phi_i$ and the mean phase $\overline{\phi}$ (see Supplementary Information Text S6). The top, middle, and bottom rows show the results in DD after T22, T24, and T26, respectively.
